# Functional Analysis of Late-Onset Alzheimer’s Disease Risk Genes in *Caenorhabditis elegans* Identifies Regulators of Neuronal Aging

**DOI:** 10.64898/2026.03.26.714531

**Authors:** Swapnil G Waghmare, Meera M Krishna, Emily C Maccoux, Ariel L Franitza, Brian A Link, Lezi E

**Affiliations:** Department of Cell Biology, Neurobiology and Anatomy, Medical College of Wisconsin, 8701 W Watertown Plank Road, Milwaukee, WI 53226, United States of America; Neuroscience Research Center, Medical College of Wisconsin, 8701 W Watertown Plank Road, Milwaukee, WI 53226, United States of America

**Keywords:** Aging, Alzheimer’s disease, LOAD, neurodegeneration, amyloid β, *Caenorhabditis elegans*, genetics

## Abstract

**Background:** Genome-wide studies in late-onset Alzheimer’s disease (LOAD) have uncovered many risk loci, yet identifying the causal genes and clarifying how these genetic signals connect to molecular and cellular mechanisms relevant to AD pathogenesis *in vivo* remains challenging.

**Methods:** Using *Caenorhabditis elegans* as a model to identify LOAD-associated genes that drive neurodegenerative processes, we focused on 14 understudied genes and their homologs: *ABI3/abi-1*, *B4GALT3/bre-4*, *CCDC6/T09B9.4*, *CLPTM1* (two homologs *C36B7.6* and *R166.2*), *CNN2/cpn-2*, *DMWD/wdr-20*, *ECHDC3/ech-2*, *MADD/aex-3*, *NCK2/nck-1*, *RABEP1/rabn-5*, *RIN3/rin-1*, *SLC39A13/zipt-13*, *TRAM1/tram-1*, and *USP6NL/tbc-17*. We knocked down these genes by RNAi and quantified lifespan, aging-associated degeneration of two neuron classes, PVD and PLM, and associative learning and short-term memory.

**Results:** Lifespan was unaffected by most knockdowns, and only *nck-1* and *tbc-17* shortened lifespan. Across neuronal assays, multiple homologs modulated aging with clear neuron-class selectivity. Knockdown of *aex-3*, *C36B7.6*, *cpn-2*, *ech-2*, *rabn-5*, *rin-1*, *T09B9.4*, and *zipt-13* attenuated late-life PVD degeneration, whereas *R166.2* and *tram-1* accelerated early PVD aging. Only two genes affected PLM aging: *R166.2* knockdown exacerbated degeneration, while *tbc-17* knockdown attenuated it despite its lifespan-shortening effect. In PLM neurons, *tbc-17* knockdown, targeting a Rab GTPase-activating protein, also preserved mitochondrial architecture during early aging and shifted heat stress-induced mitochondrial remodeling toward a pattern consistent with improved quality control. In behavioral assays, *ech-2* knockdown, targeting an enoyl-CoA-hydratase, enhanced short-term memory during early stages of aging. To further assess how LOAD-linked genes interact with Aβ-driven neurodegeneration, we developed a model that combines the PVD aging assay with a background expressing human Aβ_1-42_ panneuronally. In this model, Aβ expression accelerated age-dependent PVD degeneration, whereas *ech-2* knockdown abolished this Aβ-induced effect.

**Conclusions:** Our findings show that conserved homologs of several understudied LOAD risk genes causally modulate neuronal aging *in vivo* in a neuron-class-selective manner, often dissociable from organismal longevity. This *C. elegans* framework translates human genetic associations into quantitative, aging-linked neuronal phenotypes, and our results further emphasize early endosomal and lipid-related processes as key pathways that warrant functional testing in neuronal aging. This study also provides a tractable platform to prioritize targets for cross-species validation and to test synergy with established LOAD risk genes.

## INTRODUCTION

Alzheimer’s disease (AD) is a progressive, age-related neurodegenerative disease and the leading cause of dementia. While amyloid β (Aβ) and tau pathologies are central features of AD, in sporadic, late-onset AD (LOAD), which is strongly linked to aging and accounts for 90% of all AD cases, the etiology remains incompletely defined. The lack of effective disease-modifying or preventive therapies makes it critical to define how aging intersects with other LOAD risk factors, particularly genetic variation, to initiate and drive neuronal dysfunction, thereby revealing targetable mechanisms.

Genome-wide association studies (GWAS) have clarified core features of LOAD genetics while revealing gaps that require functional follow-up. *APOE* ε*4* appears to be the strongest genetic risk factor so far, with risk increasing per ε*4* copy (1), while ε*2* is protective (2). The locus encodes apolipoprotein E, a mediator of lipid transport and receptor-mediated lipoprotein uptake. Yet *APOE* ε*4*’s predictive value varies by ancestry: it is strong in Caucasians but lower in African American and Hispanic populations, where elevated disease risk can occur independent of *APOE* genotype (3, 4), pointing to additional genetic contributors. Ancestry-dependent effects are also evident at other loci. For example, *TREM2*, which encodes a microglial lipid-sensing receptor that controls phagocytosis and survival, has variants such as R47H that modify microglial responses to Aβ and increase risk in Europeans but show weaker or absent effects in Asian or African cohorts (5–8). Likewise, *ABCA7*, implicated in lipid transport and phagocytosis, shows particularly strong associations in African Americans and a distinct allelic architecture in Europeans (9, 10).

Recent GWAS have identified more than 75 LOAD risk loci, and their mapped genes and regulatory targets collectively implicate innate immunity, endo-lysosomal and vesicular trafficking, lipid metabolism and bioenergetic/mitochondrial pathways (11, 12). For many of these loci, we still lack functional evidence connecting risk variants to aging-dependent neuronal dysfunction and disease mechanisms (13, 14). Because most risk variants map to non-coding regulatory regions and are expected to modulate gene expression rather than protein sequence, gene-expression perturbation provides a logical initial step for functional interrogation. Dissecting how genes at these loci contribute to aging-associated neurodegeneration in mammalian models *in vivo* is essential but such studies demand substantial time and resources. Complementary approaches that allow rapid, parallel testing are therefore needed, providing a strong rationale for scalable *in vivo* models that can assay aging across the lifespan readily, to prioritize loci and effector genes, and define disease-relevant mechanisms.

*Caenorhabditis elegans* provides such a platform, with broad conservation of neurodegeneration-relevant molecular pathways, rapid genetics, and a two to three week lifespan that allows assays sampled across all aging stages and higher throughput *in vivo* testing, relative to mammalian models (15, 16). *C. elegans* PLM touch receptor neurons and PVD polymodal mechanosensory neurons are widely used in neurodegeneration studies. In particular, our prior work systematically characterized aging-associated morphological phenotypes in PVD neurons, including excessive/ectopic branching and progressive dendritic beading consistent with cytoskeleton disruption, and showed that these changes are accompanied by proprioceptive deficits and reduced harsh touch responses (17, 18). PLM neurites likewise develop age-dependent ectopic branching and lesions (19–21). These features mirror mammalian neuronal aging and neurodegeneration (22–25), and provide quantifiable, aging-sensitive readouts for genetic studies. Because these readouts are neuron-class-specific *in vivo*, the system enables principled tests of selective neuronal vulnerability, a defining feature of AD and other neurodegenerative diseases (26). Distinct *C. elegans* neuron classes follow different morphological and functional aging trajectories (27), also providing the internal contrast needed for such studies. In addition, transgenic models expressing human Aβ in *C. elegans* neurons have been shown to capture aspects of AD-relevant neurotoxicity with behavioral impairments, including learning/memory and locomotion defects (28–30). Importantly, many identified LOAD-associated genes have *C. elegans* homologs; this evolutionary conservation further supports using this model to probe the roles of these genes in aging-associated and Aβ-driven neurodegeneration.

Here, we use *C. elegans* genetics to link LOAD-associated genes to neuronal aging. We screened homologs of 14 understudied LOAD-associated genes by RNA interference (RNAi), and measured lifespan, PVD and PLM neuronal integrity, and associative learning and memory function in normal aging. Most genes did not affect lifespan. However, many knockdowns shifted the trajectory of neuronal aging, often in a neuron-class-selective manner. We then prioritized two genes with strong effects for preliminary mechanistic insight: *ech-2/ECHDC3*, which influenced PVD aging and memory function, and *tbc-17/USP6NL*, which affected PLM aging only. To further evaluate *ech-2*, we developed a new composite model that pairs the PVD aging readout with pan-neuronal Aβ expression (31, 32), enabling single-neuron visualization of Aβ-induced neuronal damage *in vivo*. *tbc-17* was assessed using a heat-shock assay that tracks mitochondrial remodeling in PLM neurons in response to stress. Together, our results provide causal links between conserved homologs of genes at understudied LOAD risk loci and neuronal aging *in vivo*. They also position *C. elegans* as a rapid, scalable platform to prioritize genes and interactions, define underlying mechanisms, and inform early-stage target discovery in mammalian models of LOAD and related neurodegenerative diseases.

## RESULTS

### Identification of understudied LOAD-associated genes and *C. elegans* homologs for *in vivo* functional assays

We assembled a candidate list of LOAD-associated genes from recent GWAS and meta-analyses (11, 12, 33–35) to investigate their roles in neurodegeneration *in vivo*. To prioritize genes with limited mechanistic characterization, we searched PubMed for each gene or protein name combined with the keywords *“Alzheimer’s”* and *“mice”*, *“mouse”*, *“in vivo”*, or *“in vitro”*. Genes with fewer than ten mechanistic studies in these contexts were classified as poorly characterized. From this filter, we selected 14 understudied LOAD genes corresponding to 15 *C. elegans* homologs identified using Ensembl Compara, InParanoid, Homologene, and OrthoMCL: *ABI3/abi-1*, *B4GALT3/bre-4*, *CCDC6/T09B9.4*, *CLPTM1* (two homologs, *R166.2* and *C36B7.6*), *CNN2/cpn-2*, *DMWD/wdr-20*, *ECHDC3/ech-2*, *MADD/aex-3*, *NCK2/nck-1*, *RABEP1/rabn-5*, *RIN3/rin-1*, *SLC39A13/zipt-13*, *TRAM1/tram-1*, and *USP6NL/tbc-17* (Table 1).

**Table. 1.** LOAD-associated genes and *C. elegans* homologs. Information on human LOAD-associated genes was compiled from ^a^ Chen et al. (33), ^b^ Hudgins et al. (34), ^c^ Schwartzentruber et al. (3) ^d^ Wightman et al. (12), ^e^ Zhang e al. (35). Human gene product functions were retrieved from GeneCards (36). Expression patterns in human aging and AD brains were aggregated from prior meta-analyses and public transcriptomic resources (37–42) (GSE118553, GSE44770, GSE48350, GSE53890). Gene expression pattern: ↑ upregulated, ↓ downregulated, - unchanged. Regions: # frontal lobe, Δ temporal lobe (including entorhinal cortex and hippocampus), † parietal lobe, ‡ cerebellum. *C. elegans* homologs were identified using Ensembl Compara, InParanoid, HomoloGene, and OrthoMCL databases. Neuronal expression levels for *C. elegans* genes were obtained from WormBase (CeNGEN) (43); +++ indicates ubiquitous expression across neurons, ++ indicates expression in the majority of neuron classes, and + indicates expression restricted to fewer than ∼20 neuron classes. AA, amino Acid; AD, Alzheimer’s disease; ER, endoplasmic reticulum; GABA, gamma-aminobutyric acid; GEF, guanine Nucleotide Exchange Factor; ZIP, zinc/iron-regulated transporter-like proteins.

| Human LOAD-associated genes |  |  |  | C. elegans homolog |  |  |
| --- | --- | --- | --- | --- | --- | --- |
| Gene | Conserved gene function | Expression in aging brain | Expression in AD brain | Gene | Neuronal expression | AA similarity to human |
| *ABI3*<sup>b,d,e</sup> | Regulates actin cytoskeleton dynamics | ↑<sup>#,Δ</sup> | ↑<sup>#,Δ</sup> | *abi-1* | ++ | 44% |
| *B4GALT3*<sup>b,c</sup> | Catalyzes β-1,4-galactosylation of glycoconjugates | ↓<sup>#</sup> | ↑<sup>#,Δ</sup> | *bre-4* | ++ | 69% |
| *CCDC6*<sup>b,c,d</sup> | Regulates DNA damage response | ↓<sup>#</sup> | ↓<sup>Δ</sup> | *T09B9.4* | ++ | 38% |
| *CLPTM1*<sup>a,b</sup> | Modulates GABA receptor binding | ↓<sup>#</sup> | ↓<sup>#,Δ</sup> | *R166.2* | +++ | 96% |
|  |  |  |  | *C36B7.6* | ++ | 93% |
| *CNN2*<sup>a,b</sup> | Organizes actin filaments | ↑<sup>#</sup> | ↑<sup>#,Δ</sup>, ↓<sup>Δ</sup> | *cpn-2* | + | 96% |
| *DMWD*<sup>a,b</sup> | Regulates deubiquitinase activity | ↓<sup>#</sup> | ↓<sup>#,Δ</sup>, ↑<sup>Δ</sup> | *wdr-20* | +++ | 47% |
| *ECHDC3*<sup>c,d,e</sup> | Catalyzes enoyl-CoA hydration; modulates insulin response | - | ↑<sup>#,Δ</sup> | *ech-2* | ++ | 85% |
| *MADD*<sup>d,e</sup> | Functions as Rab GEF; binds death receptors | ↓<sup>#</sup> | ↓<sup>#,Δ,†</sup>, ↑<sup>#,Δ,‡</sup> | *aex-3* | +++ | 38% |
| *NCK2*<sup>c,d</sup> | Functions as adaptor in receptor tyrosine kinase signaling | - | - | *nck-1* | +++ | 100% |
| *RABEP1*<sup>a,b,d</sup> | Mediates endosome trafficking; binds Rab regulators | ↓<sup>#,Δ</sup> | ↓<sup>#,Δ,†</sup> | *rabn-5* | +++ | 73% |
| *RIN3*<sup>b,c,d,e</sup> | Acts as Rab GEF | ↑<sup>#</sup> | ↑<sup>#</sup>, ↓<sup>Δ</sup> | *rin-1* | ++ | 27% |
| *SLC39A13*<sup>a</sup> | Imports zinc (ZIP family transporter) | - | - | *zipt-13* | ++ | 79% |
| *TRAM1*<sup>a,b</sup> | Facilitates ER translocation of secretory proteins | ↑<sup>#</sup> | ↑<sup>#,Δ</sup> | *tram-1* | +++ | 96% |
| *USP6NL*<sup>b,c,d</sup> | Acts as Rab GTPase-activating protein | ↓<sup>#</sup> | ↑<sup>Δ</sup>, ↓<sup>#</sup> | *tbc-17* | +++ | 36% |

Although functional links between these human LOAD genes and aging-related neurodegeneration or disease mechanisms remain limited, many of them show altered expression in human aging or AD brains based on recent meta-analyses and our secondary analysis of public brain transcriptomic datasets (Table 1; Table S1). The shared direction of expression change in aging and AD brains for most genes supports a potential aging-centered contribution to neurodegeneration. To further highlight the aging relevance in our model, we also found age-dependent expression changes in the corresponding *C. elegans* homologs, with a subset shifting in the same direction as human datasets (Fig. S1a).

In this study, we selected feeding RNAi to achieve submaximal knockdown that better approximates potential modest gene expression perturbations expected from most human non-coding variants, while avoiding lethality or developmental confounds common with null alleles of essential genes. Importantly, feeding RNAi supports either lifelong exposure or adulthood-specific knockdown by initiating RNAi after animals reach young adulthood, allowing us to isolate aging effects from development. To address the relative refractoriness of *C. elegans* neurons to feeding RNAi (44) and the predominantly neuronal expression of most homologs (Table 1) (43), we performed the screen in a neuronal RNAi-sensitized background (LC108 *uIs69 [unc-119p::sid-1]*) (45). RNAi efficiency was confirmed by qPCR (Fig. S1b-c).

For the primary screen, we used lifelong RNAi and systematically targeted each homolog and evaluated three outcomes relevant to aging and neurodegeneration: (1) lifespan, (2) aging-associated structural changes in PLM and PVD neurons, and (3) olfactory associative learning and short-term memory (Fig. 1a).

**Figure 1:**
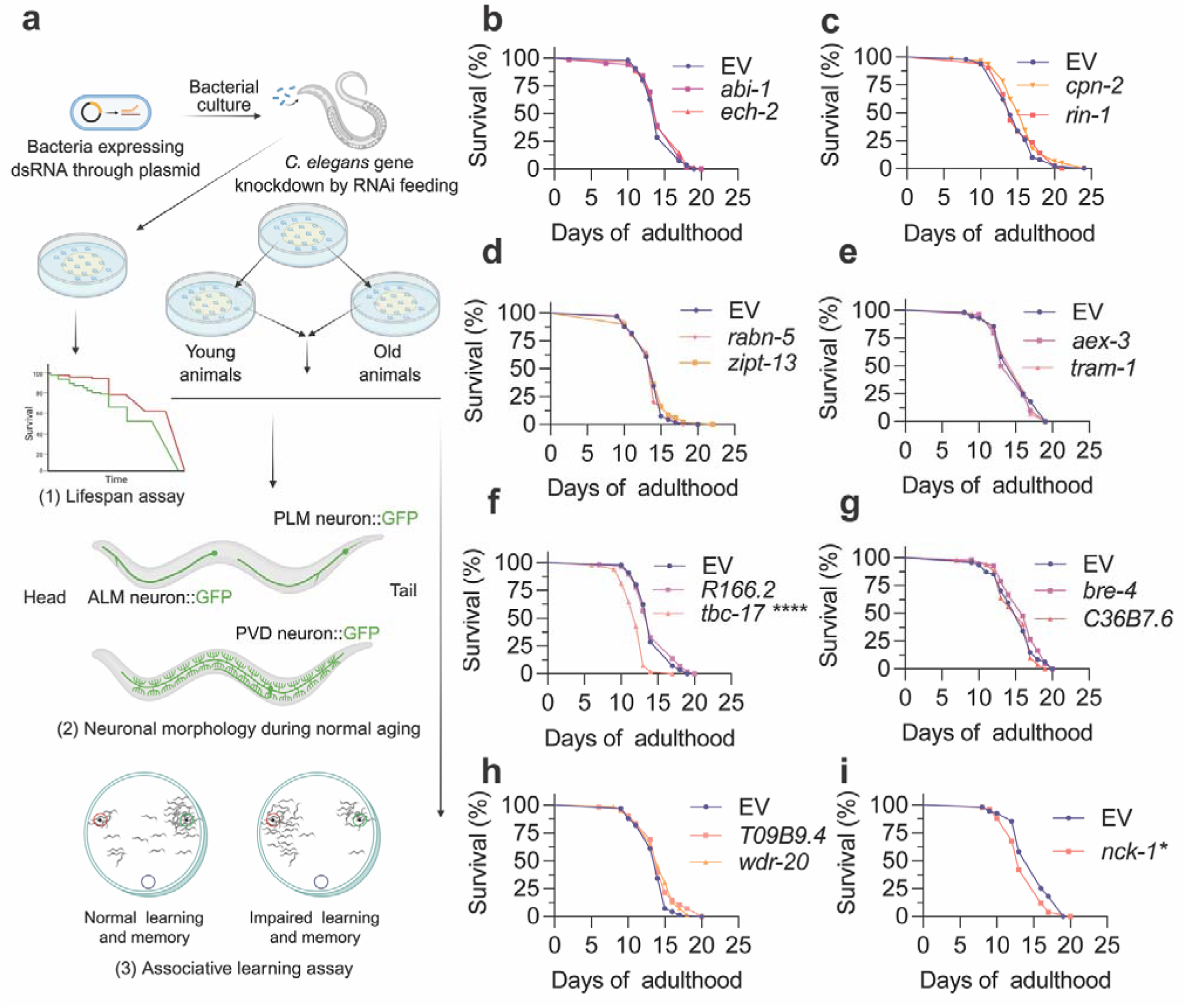
Lifelong RNAi knockdown of LOAD-associated gene homologs in *C. elegans*. **a** Schematic of the RNAi screen workflow. **b–i** Lifespan of *C. elegans* following lifelong RNAi knockdown of prioritized *C. elegans* homologs of LOAD-associated genes. Lifespan data were analyzed using Kaplan–Meier survival analysis with log-rank (Mantel–Cox) tests. EV, animals fed on *E. coli* HT115 (DE3) bacteria with empty vector L4440, served as controls. See Table S2 for additional replicates. Experiments performed without FUDR. *\* p < 0.05*, *\*\*\*\* p < 0.0001*.

### Lifespan is largely unaffected by LOAD gene homolog knockdowns

Given the strong link between age and LOAD risk, we first asked whether modulating candidate homologs alters organismal aging. Each target gene was knocked down individually in age-synchronized cohorts, with a matched empty vector (EV) as control (Fig. 1; Table S2). Survival was scored daily at 20 °C without fluorodeoxyuridine (FUDR), to avoid confounding effects on aging (46). Two knockdowns shortened lifespan: *tbc-17*, encoding a TBC-domain Rab GTPase-activating protein (GAP) predicted to act in the Rab5 pathway (47), and *nck-1*, encoding an Src homology2/3-containing adaptor protein (Fig. 1f, 1i). These effects suggest essential roles for these genes consistent with their predicted functions in endocytic trafficking and cytoskeleton-regulating signaling (47, 48). By contrast, for all other tested targets, lifespan did not differ from modifiers (Fig. 1b-h). Therefore, downstream neuronal morphology and learning assays can be interpreted without a global frailty confound for most genes.

### LOAD gene homologs modulate neuronal aging patterns in a neuron-class-selective manner

To capture progressive decline in neuronal integrity during aging, we quantified structural changes in two *C. elegans* mechanosensory neuron types, PVD and PLM. PVD neurons are a bilateral pair, and each cell body extends a single axon to the ventral nerve cord and highly elaborate dendritic arbors (Fig. 2a) that enable polymodal sensation, including harsh touch, cold, and proprioceptive/postural cues (49, 50). During normal aging, PVD dendrites progressively develop bead or buddle-like structures enriched with fragmented and discontinuous microtubules (Fig. 2a) (17, 51), closely resembling degenerative changes observed in mammalian neurons (22–24). PVD axons and cell bodies do not typically display degenerative phenotypes within the same time frame (17). PLM is a bilateral pair of gentle-touch receptor neurons; each neuron extends one long anterior primary neurite toward the mid-body and a short posterior neurite (Fig. 3a) (52). With age, PLM neurites acquire structural abnormalities such as sharp bends and ectopic branches (Fig. 3a), also reflecting cytoskeletal decline (19–21, 53). Both PLM ectopic branching/sharp bends and PVD dendritic beading are considered intrinsic features of aging, given that reducing insulin/IGF signaling through *daf-2* loss-of-function, which slows *C. elegans* aging (54), significantly delays the onset of these neuronal morphological changes (17, 21).

**Figure 2:**
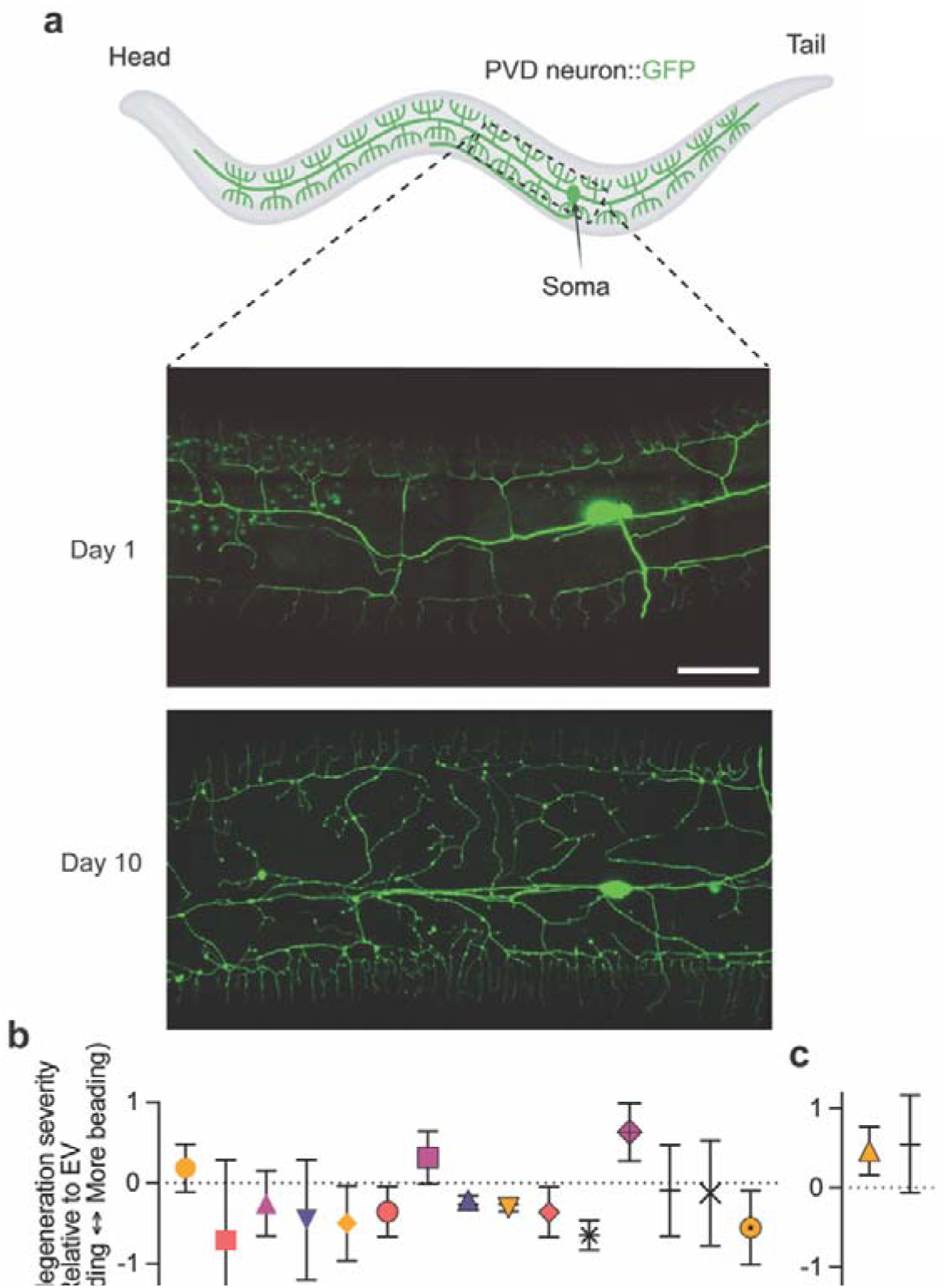
RNAi knockdown of LOAD-associated gene homologs modulates aging-associated dendritic degeneration in PVD neurons. **a** Representative confocal images of PVD mechanosensory neurons, showing its complex dendritic architecture and typical age-related dendritic beading in control animals (in neuronal RNAi sensitized background). Mild autofluorescence of gut granules, a normal physiological feature in *C. elegans*, appears as small punctate structures in the background. Scale bar = 20 µm. **b** Quantification of dendritic beading severity in PVD neurons at Day 14 following RNAi knockdown of LOAD gene homologs prioritized in this study. **c** Quantification of dendritic beading severity in PVD neurons at Day 11 following RNAi knockdown of *R166.2* and *tram-1*. PVD beading severity was scored in four categories (normal, mild, moderate, severe) based on bead number across the dendritic tree and assigned ordinal values 0–3 (see Methods). For each RNAi condition and replicate, a _Δ_-Severity value was calculated as the average severity score in the LOAD gene RNAi group minus that of its paired EV control to visualize the direction and magnitude of morphological change; _Δ_-Severity was used only for descriptive plotting. Statistical inference used the 4-level categorical data and a stratified Cochran–Mantel–Haenszel (CMH) trend test across

**Figure 3:**
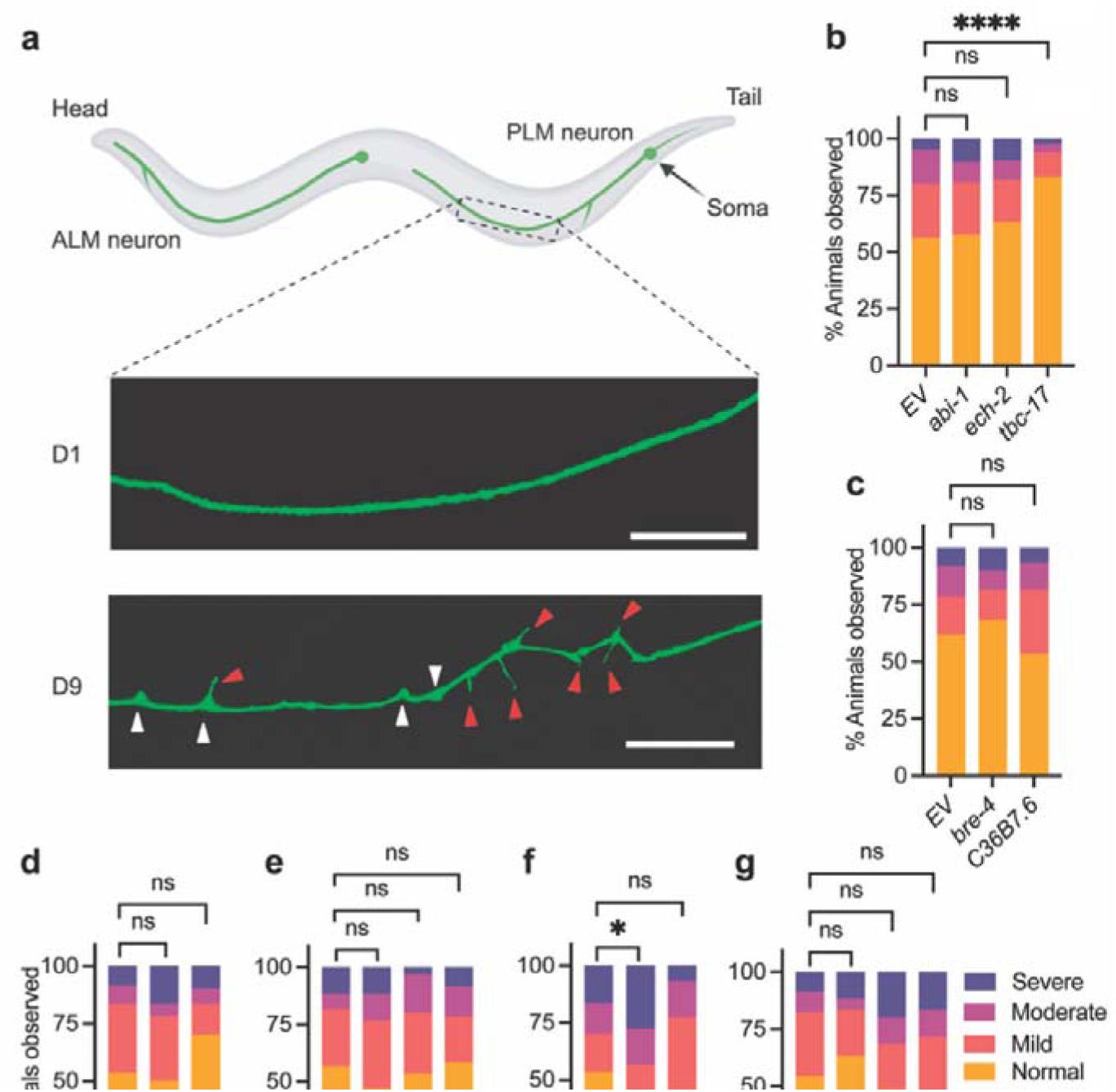
Effects of LOAD gene homologs on aging-associated morphological changes in PLM neurons. **a** Representative confocal images of PLM touch receptor neurons showing normal neurite structure and age-associated morphological changes, including ectopic branching (indicated by red arrowheads), sharp bends/kinks (indicated by white arrowheads). Scale bar = 20 µm. **b–g** Quantification of ectopic branching severity in PLM neurons at Day 9 following RNAi knockdown of LOAD gene homologs prioritized in this study. Data represent pooled results from independent biological replicates (n ≥ 60 animals per condition). Statistical comparisons were performed using

Multiple knockdowns shifted PVD aging trajectory (Fig. 2; Table S3). At Day 14 of adulthood, which is considered advanced age for *C. elegans*, knockdown of *aex-3/MADD*, *C36B7.6/CLPTM1*, *cpn-2/CNN2*, *ech-2/ECHDC3*, *rabn-5/RABEP1*, *rin-1/RIN3*, *T09B9.4/CCDC6*, and *zipt-13/SLC39A13* each reduced dendritic beading relative to empty-vector controls (Fig. 2b). Many of these protective effects were not observed at Day 11 (Fig. S2b), indicating that their effects emerge predominantly in late life, in line with the late-onset manifestation of LOAD. Two targets showed an earlier shift: *R166.2/CLPTM1* and *tram-1/TRAM1* RNAi animals did not differ from controls at Day 14 but exhibited increased PVD beading at Day 11, and neither had altered lifespan (Fig. 1e-f, Fig. 2b-c), suggesting an acceleration of aging onset with similar late-life severity. In comparison, *nck-1/NCK2* and *tbc-17/USP6NL* RNAi exacerbated PVD beading at Day 14 (Fig. 2b) but also reduced lifespan (Fig. 1f, 1i); therefore, their neuronal effects may be confounded by organismal frailty. Importantly, at Day 3, PVD morphology in RNAi conditions that later worsened PVD aging (*R166.2*, *tram-1*, and *tbc-17*) was indistinguishable from control (Fig. S2), indicating that the phenotypes reflect aging effects rather than developmental defects.

Only two knockdowns altered the aging trajectory of PLM. *R166.2/CLPTM1* RNAi increased aging-associated ectopic branching, whereas *tbc-17/USP6NL* RNAi reduced ectopic branching at Day 9 (Fig. 3b, f). Interestingly, the PLM-protective effect of *tbc-17* RNAi contrasts with its lifespan-shortening effect (Fig. 1f), indicating a partial uncoupling between organismal longevity and neuronal aging mechanisms. No targets affected frequency of neurite sharp bends, another previously described PLM aging phenotype (19, 20), and PLM morphology at Day 3 was also unaffected under all RNAi conditions (Fig. S3).

Surprisingly, two *C. elegans CLPTM1*-related homologs, *C36B7.6* and *R166.2*, showed dissociable effects on neuronal aging. *R166.2* RNAi accelerated aging in both PVD and PLM (Fig. 2c, Fig. 3f), aligning with human datasets where *CLPTM1* is downregulated in aging and AD (Table 1). In contrast, *C36B7.6* RNAi reduced PVD beading and had no detectable effect in PLM (Fig. 2b, Fig. 3c). Notably, while both paralogs are neuron-enriched, their expression patterns differ across neuron classes in *C. elegans* (Fig. S4). Taken with the paralog-specific outcomes, this points to potential cell-type specific roles for CLPTM1-linked pathways relevant to late-life neuronal maintenance, and the isoform diversity of human CLPTM1 makes selective engagement by cell type plausible, although untested.

Together, these results reveal that *C. elegans* homologs of many LOAD-associated genes are involved in regulating neuronal aging, with clear neuron-class selectivity. These findings also highlight PVD aging phenotypes as a sensitive readout of how gene expression changes impact neuronal integrity across adulthood.

### Selected LOAD gene homologs modulate short-term memory

We next asked whether *C. elegans* homologs of LOAD-associated genes influence cognitive-like functions, using a chemotaxis-based associative learning and short-term memory assay. We employed the well-established isoamyl alcohol conditioning paradigm (34, 55–57), where animals were trained to associate the attractive odor isoamyl alcohol with starvation, and chemotaxis was quantified immediately after training (0 hours post-conditioning, 0hr PC) to evaluate learning, and again one hour later (1hr PC) to assess short-term memory-like performance. In this process, olfactory chemotaxis is driven by amphid sensory neurons such as AWC, while interneurons including cholinergic AIY and AIA, and glutamatergic RIA contribute to learning and memory formation (58–60). Given the predominant vulnerability of cholinergic and glutamatergic systems in AD patients (61, 62), this assay tests LOAD-homolog effects in circuits directly implicated in the disease. After training, animals with normal learning and memory avoid the isoamyl alcohol source, whereas animals with deficits remain attracted (Fig. 4a).

**Figure 4:**
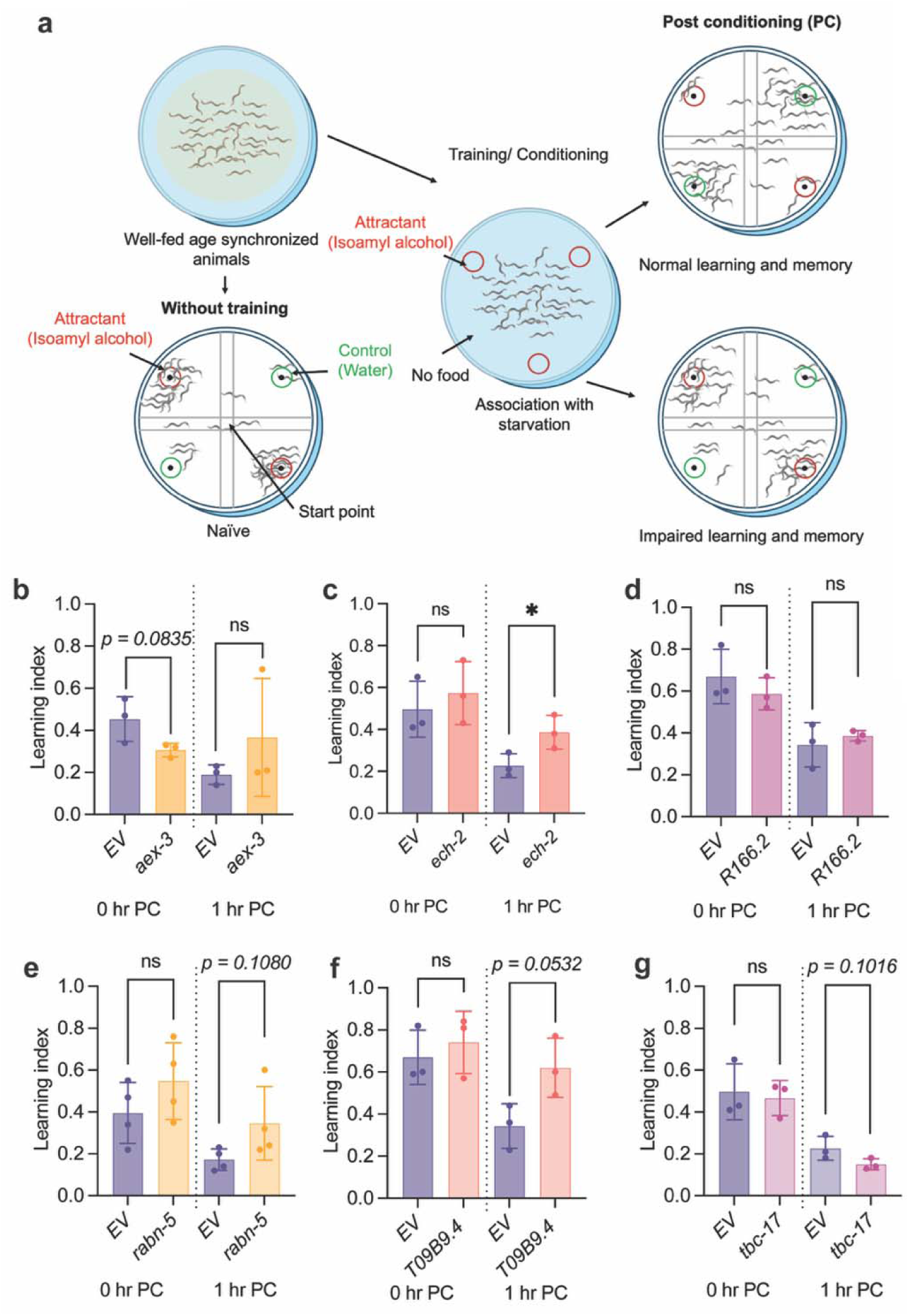
Selected LOAD gene homologs modulate memory-like behavior in *C. elegans*. **a** Schematic of the isoamyl alcohol associative learning assay. Animals are trained to associate the odorant isoamyl alcohol with starvation. Chemotaxis index is measured immediately after conditioning (0hr PC) and one hour later (1hr PC) to assess learning and short-term memory-like performance. **b–g** Quantification of chemotaxis behavior at Day 5 following lifelong RNAi knockdown of selected LOAD gene homologs. Each data point represents an independent replicate with n > 100 animals per replicate per condition. Statistical comparisons were performed using unpaired two-tailed *t*-tests. Data are presented as mean ± SD. Experiments performed with FUDR. EV, control animals. \**p < 0.05*.

Day 5 mid-adulthood animals subjected to lifelong RNAi were tested. This time point was chosen to capture early signs of age-related decline in learning and memory in control animals (Fig. S5a-b) while minimizing late-life frailty confounding chemotaxis readouts, including slowed movement and age-related muscle loss. We prioritized six LOAD gene homologs based on their effects on PLM and PVD neuronal aging described above (Figs. 2–3): *aex-3/MADD*, *ech-2/ECHDC3*, *R166.2/CLPTM1*, *rabn-5/RABEP1*, *T09B9.4/CCDC6*, and *tbc-17/USP6NL*.

We found that *ech-2* knockdown significantly enhanced memory-like behavior (Fig. 4c). *T09B9.4* RNAi animals also showed a marginal improvement in memory (Fig. 4f), and *rabn-5* knockdown showed a directionally similar but non-significant change (Fig. 4e). Learning acquisition at 0hr PC was comparable to controls for all three groups (Fig. 4). Notably, these same knockdowns also attenuated late-life PVD aging (Fig. 2b). Because the olfactory memory circuit is anatomically distinct from PVD, this pattern indicates that certain LOAD homolog perturbations influence neuronal integrity across multiple neuron populations.

*tbc-17* or *R166.2* knockdown did not alter learning acquisition or 1 hr PC performance (Fig. 4d, g). *aex-3* knockdown showed a small, non-significant reduction in learning at 0hr PC without affecting short-term memory (Fig. 4b). Naïve chemotaxis (without training/conditioning) was comparable between all RNAi groups and EV controls, including *aex-3* RNAi (Fig. S5c-h), suggesting the observed differences reflect associative acquisition rather than altered primary trend and protection of PVD aging with *aex-3* RNAi (Fig. 2b) points to neuronal population-specific impacts that yield divergent functional and behavioral consequences during aging.

### Neuron-class selective impacts of *tbc-17/USP6NL* during aging are linked to mitochondrial network remodeling

We next prioritized *tbc-17/USP6NL* for further follow-up because its RNAi produced a paradoxical profile, with shortened lifespan (Fig. 1f) and exacerbated PVD aging (Fig. 2b), yet reduced age-associated PLM ectopic branching (Fig. 3b), motivating investigation of the underlying pathway. *tbc-17* encodes a TBC-domain Rab GAP predicted to act in the Rab5 pathway (47). While the overall protein similarity between *C. elegans* TBC-17 and human USP6NL is modest (∼36%, Table 1), the TBC/Rab GAP domain is highly conserved, supporting functional inference across species (Fig. S6a).

To further confirm the role of *tbc-17* in PLM aging, we constructed a second, independent RNAi clone; knockdown with this clone reproduced the phenotype from initial screen (Fig. 3b; Fig. S6b). We then performed adulthood-specific RNAi to ensure that the observed effects do not arise from altered embryonic or larval development. Knockdown was initiated at L4 stage (last larval stage, early young adulthood), and animals were phenotyped at Day 9. This regimen produced the attenuation of PLM ectopic branching similar to that seen with lifelong RNAi (Fig. 5a; Fig. 3b). Lifespan in adulthood-specific RNAi animals was comparable to controls, indicating that the shortening seen with lifelong *tbc-17* knockdown likely reflects developmental effects (Fig. 1f; Fig. S6c). Conversely, overexpressing *tbc-17* under its endogenous promoter increased PLM ectopic branching in Day 3 young adults, despite increased lifespan (Fig. 5b; Fig. S6d-f). To determine whether this morphological change impairs neuronal function, we measured animal’s responses to gentle touch, a modality sensed by the PLM neurons (63, 64). In wild-type animals, sensitivity to gentle touch declined significantly with age (Fig. S6g), but *tbc-17* overexpression did not alter touch responsiveness at Day 3 (Fig. S6h). The absence of an early-life behavioral deficit despite increased ectopic branching in PLM may reflect compensatory mechanisms within the mechanosensory circuit or a threshold requirement before neuronal structural changes are sufficient to disrupt signaling.

**Figure 5:**
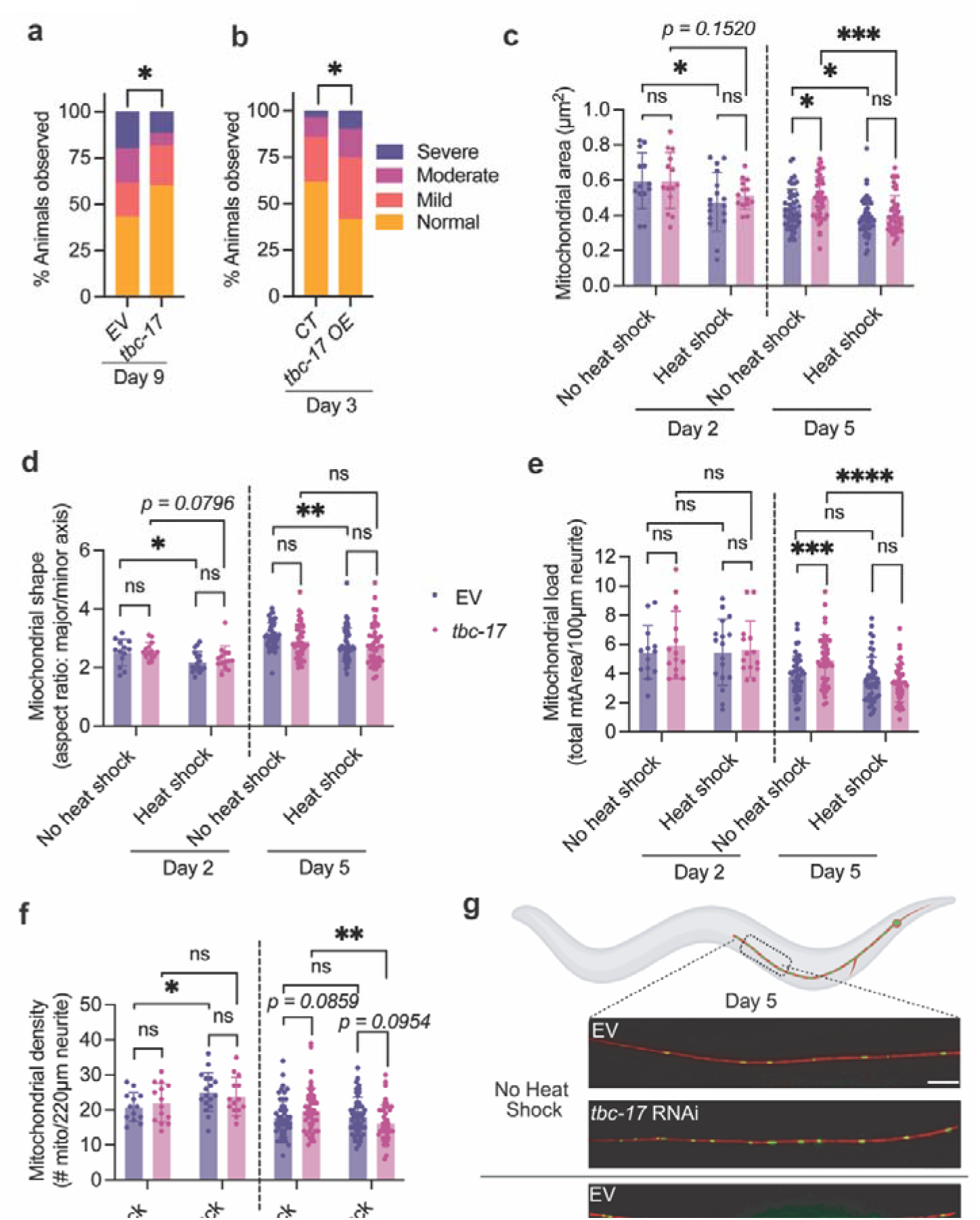
Neuron-class selective impacts of *tbc-17/USP6NL* during aging are linked to mitochondrial dynamics. **a** Quantification of ectopic branching severity in PLM neurons at Day 9 following adulthood RNAi knockdown of *tbc-17* (n = 60) and EV (n = 60). Experiments performed with FUDR. **b** Quantification of ectopic branching in PLM neurons of control (CT, ELZ238) (n = 139) and *tbc-17* overexpression animals (*tbc-17* OE, ELZ279) (n = 54) at Day 3. See Figuire S6 for an additional line. Animals were fed on standard *E. coli* OP50. Experiments performed with FUDR. **c–f** Quantification of mitochondrial morphology in PLM neurons of Day 2 and Day 5 adults following lifelong *tbc-17* knockdown, under basal and heat stress conditions. Each data point represents an individual animal. Experiments performed without FUDR. **g** Representative images of PLM neurons under control and heat shock conditions in the EV and *tbc-17* lifelong RNAi groups at Day 5. Data are presented as mean ± SD. PLM neurite is labeled with RFP, and PLM mitochondria are labeled with GFP. Scale bar = 10 µm. In **a** and **b,** statistical comparisons used Mann–Whitney tests. Data in **c–f** were

We next explored mechanisms by which *tbc-17* regulates PLM aging. Neuronal mitochondrial dysfunction is a key feature of AD pathogenesis, compromising synaptic maintenance, axonal transport, and neuronal survival (65, 66). Neuronal mitochondria undergo age-related changes in size, shape and spatial distribution (67–69); accordingly, quantitative metrics of these features are commonly used as readouts that can indicate changes in fission–fusion balance, biogenesis, or clearance when interpreted with appropriate controls. In *C. elegans*, mitochondria also accumulate abnormally at PLM ectopic branching sites (20), linking neuronal structural integrity to mitochondrial homeostasis. We therefore used a reporter strain (*jsIs609 (mec-7p::mitoGFP)*) validated for visualizing mitochondrial dynamics in *C. elegans* mechanosensory neurons (68, 70), to measure four mitochondrial metrics in PLM neurons of control and *tbc-17* lifelong RNAi animals. Mitochondrial area served as a proxy for size and was calculated as the average area per mitochondrial particle in each neuron. Mitochondrial shape was estimated by aspect ratio, defined as major axis divided by minor axis, where higher values indicate more elongation. Mitochondrial load captured total content as the summed fluorescence signal from all particles, and mitochondrial density was the number of particles per fixed length of PLM neurite.

We first assessed baseline mitochondrial morphology at Day 2 of adulthood and at Day 5, an early aging time point (Fig. 5; Fig. S6). At Day 2, mitochondrial features were comparable between control and *tbc-17* RNAi animals across all parameters. By Day 5, controls shifted toward smaller, more elongated mitochondria (higher aspect ratio) with reduced content (lower total load), compared with Day 2 (Fig. S6i-k). In *tbc-17* RNAi animals, aspect ratio did not increase significantly with age and total load at Day 5 was higher than in controls, indicating fewer age-linked changes than in EV (Fig. S6j-k). These baseline data indicate that reducing *tbc-17* limits an early-aging shift in mitochondrial morphology. Notably, at Day 5, qPCR showed no change in *drp-1* (dynamin-related protein 1, mitochondrial fission) or *fzo-1* (mitofusin ortholog, mitochondrial fusion) with *tbc-17* RNAi (Fig. S6m-n), indicating that the differences in PLM mitochondrial morphology are likely not explained by altered transcription of core fission/fusion genes.

We then tested whether *tbc-17* knockdown modulates the mitochondrial stress response. Day 2 and Day 5 animals were exposed to acute heat stress (37 °C, 1 hr), and recovered for 4 hr at 20 °C before imaging. This paradigm is widely used in *C. elegans* to induce rapid remodeling of mitochondrial networks, triggering fragmentation and engaging turnover pathways within hours (71–73). In mammalian neurons, acute thermal or oxidative challenges similarly trigger rapid shifts in mitochondrial dynamics coupled to quality-control mechanisms such as mitophagy (74), indicating a conserved response.

At Day 2, control animals responded to heat shock with robust mitochondrial fragmentation, indicated by decreased average area and aspect ratio along with increased mitochondrial density (Fig. 5c-f). Under the same conditions, *tbc-17* RNAi animals exhibited minimal mitochondrial morphological change, with only a trend toward decreased aspect ratio, indicating relative resistance to heat shock-evoked fragmentation-biased reorganization of the mitochondrial network or faster recovery within the 4-hr window (Fig. 5c-f). Although the functional benefit of this attenuation remains to be tested, it suggests less mitochondrial network disruption than in controls. At Day 5, control animals had smaller, rounder mitochondria after heat shock without net content change (Fig. 5c-f). In contrast, *tbc-17* RNAi animals showed a significant reduction in mitochondrial size, total load, and density following heat stress, without significant changes in aspect ratio (Fig. 5c-f), a response most consistent with engagement of quality-control pathways that remove damaged mitochondria rather than simple accumulation of rounded fragments. Thus, during early aging, reducing *tbc-17* appears to elicit earlier coupling of stress-induced mitochondrial reorganization to pathways for mitochondrial quality control.

Together, these findings indicate that reduced *tbc-17* expression limits early-aging neuronal mitochondrial changes and promotes stress-responsive remodeling of the mitochondrial network, a pattern that may support healthier neuronal networks.

### *ech-2/ECHDC3* knockdown attenuates Aβ-associated neurodegeneration

Whether and how recently identified LOAD-associated genes modulate Aβ-driven neurodegeneration with age is unclear. To probe this, we combined our PVD aging assay with the transgenic strain, *gnaIs2 [myo-2p::YFP + unc-119p::A*β*1-42]*, which expresses human Aβ_1-42_ pan-neuronally in *C. elegans* at relatively low levels. The *gnaIs2* Aβ model exhibits age-progressive Aβ aggregation without markedly shortening lifespan and has been reported to develop progressive impairments in mitochondrial function, olfactory chemotaxis, and locomotion (31, 32, 75). We selected *gnaIs2* over higher-expression Aβ models because it avoids severe developmental delays and lifespan reduction, enabling analysis of late-age phenotypes. To obtain a neuron-specific readout of degeneration driven by Aβ, we crossed *gnaIs2* into a PVD-specific GFP reporter line in a neuronal RNAi-sensitized background (*unc-119p::sid-1*) (Fig. 6a). This approach addresses a common limitation of prior *C. elegans* Aβ studies that relied on locomotion-based assays, which are susceptible to age-related muscle decline (76). We first compared the Aβ x PVD strain with the non-Aβ control across age. PVD beading was indistinguishable at Day 1 and Day 3, became of PVD aging by Aβ overexpression was also observed without the RNAi-sensitized background (Fig. S7c). Additionally, a strain overexpressing high levels of human Aβ in muscles that exhibits strong early toxicity (77, 78) displayed marked PVD beading at Day 1 (Fig. S7d), further confirming the sensitivity of the PVD aging model to Aβ-driven neurodegeneration *in vivo*.

**Figure 6:**
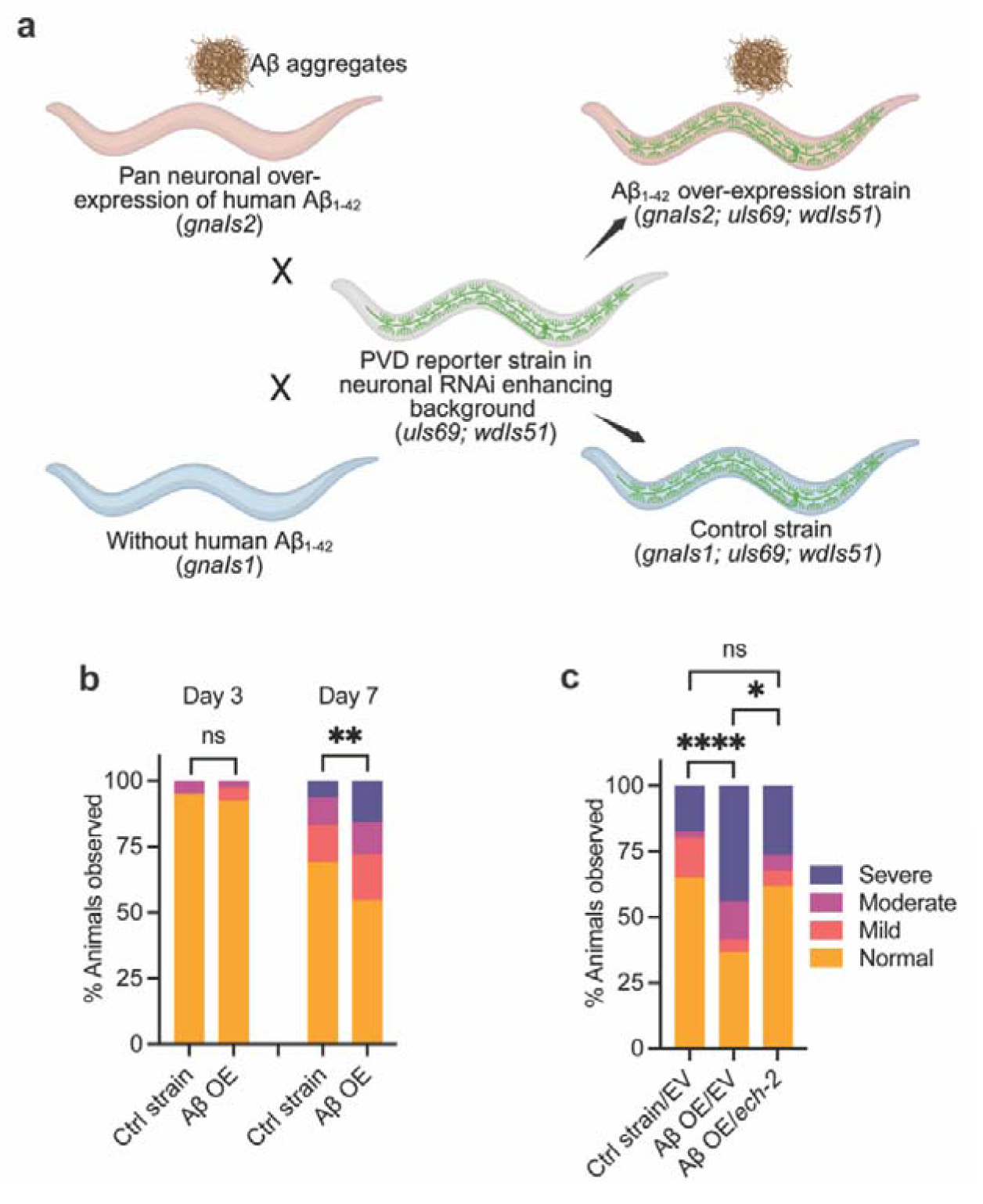
e*c*h*-2/ECHDC3* knockdown attenuates A_β_-associated neurodegeneration. **a** Schematic of strains used to visualize A_β_-associated changes in PVD neurons. **b** Quantification of aging-associated dendritic beading in PVD neurons in control animals at Day 3 (ELZ273, n = 40), in A_β_ overexpression (OE) animals at Day 3 (ELZ266, n = 39), in control animals at Day 7 (ELZ273, n = 94), and in A_β_ OE animals at Day 7 (ELZ266, n = 75). **c** Quantification of dendritic beading severity in PVD neurons at Day 9 following RNAi knockdown of *ech-2* in the A_β_-overexpressing background. Control strain ELZ273 treated with EV, n = 63; A_β_ OE strain ELZ266 treated with EV, n = 75; A_β_ OE strain ELZ266 treated with *ech-2* RNAi, n = 59. Statistical comparisons were performed using the stratified CMH trend test. See Figure S7 for additional replicates. Experiments performed with FUDR. EV, control animals fed with empty L4440 RNAi vector. *\*p < 0.05, **p < 0.01, ****p < 0.0001, ns: not significant*.

We next asked whether *ech-2/ECHDC3* modulates Aβ-associated degeneration. We prioritized *ech-2* because its knockdown preserved PVD integrity and improved short-term memory retention during aging (Fig. 2b; Fig. 4c). Additionally, it shares high homology with human *ECHDC3* (Table 1), which encodes a mitochondrial protein with an enoyl-CoA-hydratase like domain predicted to participate in fatty-acid β-oxidation. We found that lifelong *ech-2* RNAi significantly reduced Aβ-induced PVD beading in late mid-adulthood, restored to non-Aβ control levels (Fig. 6c; Fig. S7b), suggesting a functional interaction where *ech-2* reduction engages resilience mechanisms in the presence of Aβ.

To explore whether *ech-2* acts through known mechanisms underlying PVD aging, we examined NLP-29 (neuropeptide-like protein 29). NLP-29 is an epidermal antimicrobial peptide that increases with age in the absence of infections and drives PVD dendritic degeneration via neuronal NPR-12/GPCR and downstream autophagy activation (17). *ech-2* knockdown did not significantly alter *nlp-29* expression (Fig. S7e), suggesting that *ech-2*’s effect on neuronal aging and Aβ-associated degeneration are unlikely to be mediated by the NLP-29/NPR-12 pathway and pointing instead to alternative mechanisms.

## DISCUSSION

Genome-wide studies in LOAD have identified many risk loci, yet their links to targetable molecular and cellular mechanisms relevant to AD pathogenesis remain limited. In this study, we address this gap by investigating the conserved *C. elegans* homologs of understudied LOAD-associated genes and measuring effects on neuronal health across the lifespan with cell-type resolved imaging and circuit-level learning and memory behavior. Across these measures, many LOAD gene homolog knockdowns show neuron-class selective effects rather than pan-neuronal changes. We also establish an aging-relevant model that quantifies Aβ-induced damage at single-neuron resolution *in vivo*, providing a new tool to test how modulating LOAD risk genes shifts susceptibility to Aβ while making selective neuronal vulnerability, a key feature of AD, measurable. Overall, our data emphasize testable roles for Rab5-centered endosomal control (*RIN3*, *RABEP1*, *USP6NL*) and mitochondrial lipid handling (*ECHDC3*) in neuronal maintenance across aging and in AD-relevant contexts. By translating human genetic associations into quantitative *in vivo* phenotypes, this work provides a practical route to prioritize LOAD risk genes, test gene-gene interactions, and focus on cross-species validation.

*ECHDC3* encodes a mitochondrial protein with an enoyl-CoA-hydratase like domain predicted to participate in fatty-acid β-oxidation. Human studies report higher *ECHDC3* expression in AD brains than in control subjects (39) and genetic variation at this locus (e.g., rs11257242-C) is associated with hippocampal structure and cognitive alterations (79–81). Evidence from the same enzyme class shows that disrupting enzyme activity can degrade neuronal health. For example, mutations in ECHS1, encoding the mitochondrial short-chain enoyl-CoA hydratase, cause Leigh syndrome with encephalopathic presentations and are mechanistically linked to accumulation of reactive intermediates, glutathione depletion and impaired mitochondrial respiration (82, 83). For long-chain fatty acids, the enoyl-CoA hydratase step is carried out by the mitochondrial trifunctional protein (HADHA/HADHB). Defects in this complex reduce long-chain β-oxidation and are linked to peripheral neuropathy consistent with bioenergetic stress in neural tissues (84). Beyond energy failure and redox imbalance, these hydratase defects also disrupt lipid metabolism, including lipid-droplet accumulation and shifts in sphingolipid pathways (85, 86). Our *in vivo* studies show that lowering the conserved *ech-2* attenuates aging-associated and Aβ-induced neurodegeneration without extending lifespan, indicating a potential neuron-intrinsic lipid-handling axis separable from organismal aging. Because acyl-CoA pools feed lipid synthesis, remodeling, and oxidation, small flux shifts can alter membrane lipids, including phospholipids, sphingolipids, and cholesterol esters, which are broadly altered in AD (87). Such shifts can potentially modify Aβ membrane association, oligomer stability, and endocytic routing, thereby altering toxicity (87). Together, these links warrant testing whether *ECHDC3*-linked acyl-CoA metabolism could modulate neuronal Aβ susceptibility by shaping membrane lipid composition.

Maintaining mitochondrial integrity is a central task in aging neurons. Early endosomal Rab5 machinery has been shown to contribute to this task. Under mitochondrial stress, Rab5 effectors are recruited to damaged mitochondria, increasing endosome-mitochondria contacts and enabling endosomal sorting complex required for transport (ESCRT)-linked sequestration and lysosomal disposal in a Drp1-independent manner that complements canonical mitophagy (88). In this context, *tbc-17* knockdown preserves PLM neuronal integrity with age and maintains PLM mitochondrial architecture in aging and after heat stress, with a pattern most consistent with rapid pruning and clearance of damaged mitochondria rather than accumulation of fragments. TBC-17 carries a TBC/Rab-GAP domain highly conserved with the human ortholog *USP6NL* (previously known as *RN-tre*), and the related TBC-2 acts as a Rab5 GAP *in vivo* (89), supporting assignment of TBC-17 to the Rab5 pathway. In mammalian cells, USP6NL localizes to Rab5-positive endosomes and regulates Rab5-dependent cargo trafficking, although biochemical work suggests that RN-tre can also act as a GAP for other Rabs such as Rab41 (90, 91). Functionally, *USP6NL* modulates receptor endocytosis and trafficking; in cancer cells it slows epidermal growth factor receptor internalization to sustain PI3K/AKT signaling and stabilizes GLUT1, glucose transporter type 1, to support glycolytic metabolism (92–94). Taken together with our *tbc-17* findings, these properties place *tbc-17/USP6NL* in a Rab5-governed, endosome-to-mitochondrial quality-control route and support a working model in which modest GAP reduction would prolong Rab5 activity and enhance clearance of damaged mitochondria to preserve neurons independent of lifespan.

A Rab5-positive early-endosome axis appears to be a shared route influencing neuronal aging across multiple conserved LOAD gene homologs, beyond *tbc-17/USP6NL*. *rabn-5/RABEP1* encodes the Rab5 effector Rabaptin-5 and *rin-1/RIN3* a Rab5 family guanine nucleotide exchange factor (GEF) that promotes Rab5 activation and endocytic trafficking. In mammalian systems, elevated Rab5 activity enlarges early endosomes and shifts APP trafficking toward increased Aβ production, whereas moderating Rab5 reduces amyloidogenic processing of APP (95–97). Consistent with this, an AD-associated RABEP1 variant (p.R845W) hyperactivates Rab5 in HEK293T cells, enlarges early endosomes and increases Aβ42 (98). RIN3 also recruits the protein products of the LOAD risk genes, *BIN1* and *CD2AP*, to Rab5-positive endosomes in APP/PS1 mouse brains, altering APP processing and axonal transport (99). Most prior work has emphasized Rab5-linked changes in APP processing and endosome morphology, whereas how endosomal tone integrates with neuronal aging *in vivo* remains less defined. By defining convergence on a Rab5-centered route, we translate diffuse LOAD GWAS signals into prioritized targets and testable hypotheses for how endosomal trafficking supports normal neuronal aging and how its dysregulation can promote neurodegeneration.

Interpreting direction of effect for LOAD risk genes depends on dose and cell type. Our study emphasized partial knockdown of *C. elegans* homologs and did not broadly test gain of function. For example, knockdown of *abi-1*, *bre-4*, and *wdr-20* did not alter neuronal aging in our assays, yet their human homologs *ABI3*, *B4GALT3*, and *DMWD* are increased in AD cortex (Table 1), suggesting that increased dose may be the relevant manipulation to reveal phenotypes. However, bulk brain RNA from postmortem tissue provides limited guidance because it averages mixed cell types and disease stages, and blurs post-transcriptional control. Direction of effect may therefore be better inferred from cell-type-resolved readouts. Our assays already point to this need for context: several homolog knockdowns produced neuron-subtype-selective outcomes and benefits to neuronal aging without changes in lifespan. Human data also show cell-type specificity at LOAD loci. At the *USP6NL* locus, for example, the risk allele rs7912495-G is associated with higher *USP6NL* expression in microglia, while little change is detected in bulk cortex (100) (Table 1), demonstrating that tissue averages can miss cell type-restricted regulation. Accordingly, direction of effect is better established with cell type-resolved and longitudinal readouts *in vivo*. As a next step, titrated expression of human allelic variants in defined neurons or non-neuronal tissues in *C. elegans* could help clarify how expression level and cellular context shape their impact on neuronal aging.

Selective neuronal vulnerability is a defining feature of normal brain aging and many age-related neurodegenerative diseases (26). It likely reflects the intersection of neuron-intrinsic demands and extrinsic cues, yielding cell-type specific sequences of early degeneration and dysfunction. In our studies, disrupting several LOAD homologs produce neuron class-restricted effects, with PVD showing greater susceptibility. These differences are not explained by gene expression patterns alone. For example, *ech-2* knockdown attenuates PVD aging although *ech-2* is not expressed in PVD; knockdown of *T09B9.4*, the *C. elegans* homolog of human *CCDC6*, improves short-term memory despite little or no expression in AIY, AIA, and RIA, neurons that contribute to memory circuits in *C. elegans* (43, 59, 101). These non-cell-autonomous patterns indicate that the specific neuron class, its circuit connections, and its local environment, rather than expression alone, may determine where effects occur. Features such as cell surface molecular signatures, synaptic activity, and support from surrounding cells/tissues likely shape the outcome.

In line with these non-cell-autonomous patterns, PVD could be suited to study how LOAD risk genes influence selective neuronal vulnerability. PVD’s elaborate dendritic arbor imposes sustained demands on membrane delivery, cytoskeletal maintenance, and energy metabolism, which may underlie robust phenotypes in response to neuronal aging and Aβ exposure that resemble mammalian neurodegeneration. Work from our group and others has characterized these phenotypes and identified extrinsic drivers during normal aging, including skin-derived innate immune signaling and extracellular matrix remodeling (17, 18, 51, 102). For example, an age-dependent rise in the epidermal antimicrobial peptide NLP-29 may activate the conserved neuronal GPCR NPR-12 to engage autophagic machinery and drive PVD aging, while sparing PLM, which does not express the receptor. Thus, PVD provides a system to test how LOAD risk genes influence neuronal aging through non-cell-autonomous mechanisms, including signals from other neurons and from non-neural tissues, in AD-relevant contexts.

## CONCLUSIONS

In summary, we demonstrate causal effects of understudied LOAD risk gene homologs on neuronal aging and Aβ-induced neurodegeneration *in vivo*. Our findings indicate that neuronal resilience can be dissociated from organismal longevity. They also implicate two conserved routes that shape late-life neuronal health: Rab5-centered early endosomal control and lipid metabolic handling influencing Aβ susceptibility. By converting LOAD GWAS hits into mechanism-linked phenotypes, we provide an aging-relevant platform that clarifies direction of effect by cell type, prioritizes genes and contexts for mammalian validation, and supports synergy tests with established LOAD risk variants to guide new hypotheses building.

## ABBREVIATIONS

ABCA7: ATP Binding Cassette Subfamily A Member 7
ABI3: ABI Family Member 3
Aβ: Amyloid β
AD: Alzheimer’s Disease
AIA: Anterior Interneuron A
AIY: Anterior Interneuron Y
AKT: Serine/Threonine Protein Kinase
APP: Amyloid Precursor Protein
APOE: Apolipoprotein E
AWC: Amphid Wing Neuron C
B4GALT3: Beta-1,4 Galactosyltransferase 3
BIN1: Bridging Integrator 1
CD2AP: CD2 Associated Protein
CCDC6: Coiled Coil Domain Containing 6
CLPTM1: Cleft Lip And Palate Associated Transmembrane Protein 1
CNN2: Calponin 2
DMWD: Dystrophia Myotonica WD Repeat Containing Protein
ECHDC3: Enoyl CoA Hydratase Domain Containing 3
ECHS1: Enoyl CoA Hydratase Short Chain 1
ESCRT: Endosomal Sorting Complex Required For Transport
EV: Empty Vector
FUDR: Fluorodeoxyuridine
GAP: GTPase Activating Protein
GEF: Guanine Nucleotide Exchange Factor
GLUT1: Glucose Transporter 1
GPCR: G Protein Coupled Receptor
HADHA: Hydroxyacyl CoA Dehydrogenase Trifunctional Multienzyme Complex Subunit Alpha
HADHB: Hydroxyacyl CoA Dehydrogenase Trifunctional Multienzyme Complex Subunit Beta
IGF: Insulin Like Growth Factor
LOAD: Late Onset Alzheimer’s Disease
MADD: MAP Kinase Activating Death Domain
NCK2: NCK Adaptor Protein 2
NLP-29: Neuropeptide Like Protein 29
NPR-12: Neuropeptide Receptor 12
OE: Overexpression
PC: Post Conditioning
PI3K: Phosphoinositide 3 Kinase
PLM: Posterior Lateral Microtubule Neuron
PS1: Presenilin 1
PVD: Posterior Ventral Process D
RABEP1: Rabaptin RAB GTPase Binding Effector Protein 1
RIA: Ring Interneuron A
RIN3: Ras And Rab Interactor 3
RNAi: RNA Interference
SLC39A13: Solute Carrier Family 39 Member 13
TBC: Tre 2/Bub2/Cdc16 Domain
TREM2: Triggering Receptor Expressed On Myeloid Cells 2
TRAM1: Translocation Associated Membrane Protein 1
USP6NL: USP6 N Terminal Like

## ACKNOWLEDGMENTS

Some strains were provided by the CGC, which is funded by NIH Office of Research Infrastructure Programs (P40 OD010440).

## AUTHORS’ CONTRIBUTIONS

L.E conceived the project; L.E, S.G.W. and M.M.K. designed the experiments; S.G.W., M.M.K., E.C.M., A.L.F., and L.E performed the experiments and collected the data; S.G.W., M.M.K., E.C.M. and L.E analyzed and interpreted the data; S.G.W. and L.E drafted the manuscript; S.G.W., M.M.K., B.A.L, and L.E edited the manuscript; and all authors reviewed the manuscript.

## DATA AVAILABILITY

All data supporting the findings of this study are available within the paper and its Supplementary Information.

## ETHICS DECLARATIONS

### Ethics approval and Consent to participate

Not applicable. This study used *C. elegans* as a model organism and did not involve human participants, human material, or identifiable human data.

### Consent for publication

Not applicable.

## Methods

### Strains and Maintenance

All *C. elegans* strains were maintained on nematode growth medium (NGM) plates seeded with *E. coli OP50* at 20 °C following standard procedures (103). Hermaphrodite animals were used for all experiments. For age-synchronization, mid-to-late L4 larvae were selected manually and transferred to fresh NGM plates. For some assays, 100 µM 5′-fluorodeoxyuridine (FUDR; VWR, Cat# 76345-984) was added to plates to inhibit progeny production. Day 1 of adulthood was defined as 24 hours post L4. For FUDR-free conditions, animals were transferred daily until Day 5, then every other day to avoid progeny confounds. FUDR usage is noted in corresponding figure legends. Strain details are listed in Supplementary Table S4. FUDR was avoided in lifespan assays given its potential confounding effects on organismal aging (46, 104).

### RNAi

RNAi was performed using the feeding method. Bacterial clones expressing gene-specific dsRNA were obtained from the Vidal libraries and identities were confirmed by sequencing. For target genes not available in the library, we designed and generated custom RNAi clones. The full list of RNAi constructs and primers (MilliporeSigma) used for clone generation is provided in the supplementary table S5-6. *E. coli* HT115(DE3) bacteria carrying the empty L4440 vector (EV) were used as the control. RNAi plates were prepared using NGM supplemented with 1 mM isopropyl β-D-1-thiogalactopyranoside (IPTG, Sigma-Aldrich, Cat# I6758) and 25 µg/mL carbenicillin (Sigma-Aldrich, Cat# C3416). Neuronal RNAi sensitized strain was used for all experiments, unless otherwise stated. The *uIs69 [unc-119p::sid-1]* strain enables effective neuronal RNAi by expressing *sid-1* under the pan-neuronal *unc-119* promoter, thereby enhancing RNAi sensitivity in neurons (45). This strain was crossed into appropriate reporter backgrounds. For lifelong RNAi, treatment was initiated at the L4 stage of the parental generation and maintained throughout the lifespan of the F1 progeny. For adulthood-specific RNAi, treatment began at the L4 stage of the same generation and continued into adulthood. The type of RNAi timing (lifelong or adulthood-specific) used in each experiment is specified in the corresponding figure legends and methods subsections.

### Lifespan Assay

Lifespan assays were conducted at 20 °C in the absence of FUDR. Animals were monitored daily starting from Day 1 of adulthood. Animals were transferred daily until Day 5 of adulthood and then every other day thereafter to minimize confounding from progeny. Lifespan was calculated from the time animals were first placed on the assay plates until they were scored as dead. An animal was considered dead when it failed to respond to gentle prodding with a platinum wire and showed no pharyngeal pumping. Animals that died due to vulval rupture, internal hatching (bagging), or desiccation (crawling off the agar) were censored from the analysis. Each condition included 90–100 animals across three biological replicates.

### Fluorescent Microscopy and Quantification of Neuronal Aging

PLM neurons were visualized using *zdIs5* (*mec-4p::GFP*), and PVD neurons were visualized using *wdIs51* (*F49H12.4::GFP*). Animals were immobilized in 5 μM levamisole and mounted on 4% agarose pads in M9 solution. Imaging was performed using a Zeiss Axio Imager M2 compound microscope or a Leica SP8 confocal microscope with 63× oil immersion objectives. Confocal Z-stacks were acquired with 0.5–1 μm optical sections, and imaging parameters were kept constant across genotypes within each experiment. All experiments were performed in the presence of FUDR.

PLM neurite morphology was analyzed in Day 3 and Day 9 adults. Ectopic branches and sharp angular kinks disrupting neurite continuity were scored. Ectopic branching severity was categorized as follows based on the number of ectopic branches observed: 0 ectopic branches (normal), 1 (mild), 2 (moderate), and >3 (severe). PVD dendritic beading was evaluated in Day 3, Day 11 and Day 14 adults by examining the entire dendritic tree of either PVDL or PVDR. Beads were defined as bead- or bubble-like spherical enlargements with increased GFP signal. Animals exhibiting ≥ 10 beads across ≥ 5 menorahs were classified as beading-positive, indicating degenerative change (17). Degeneration severity was categorized as follows: 0–10 beads (normal), 10–20 (mild), 20–50 (moderate), and >50 (severe). Each replicate included 40-50 animals. All scoring was performed blinded to genotype and RNAi condition. To better visualize shifts in PVD beading severity and to summarize effects across independent replicates in graphs, we defined Δ-Severity relative to the paired EV control in each replicate. We assigned scores Normal = 0, Mild = 1, Moderate = 2, Severe = 3, and for each replicate and group, we computed an average severity score by weighting the animal counts in each category: average severity score = [(0 × Normal count) + (1 × Mild count) + (2 × Moderate count) + (3 × Severe count)] ÷ total animals in that group. Δ-Severity was the score in the LOAD gene RNAi group minus the score of its paired EV control within the same replicate. Negative values indicate a shift toward milder outcomes, and positive values indicate a shift toward more severe outcomes. This metric was used for visualization only in Figure 2 and Figure S2, and the mean Δ-Severity across independent replicates with SD was displayed. For statistical analysis, see below.

### Associative Learning and Short-term Memory Assay

We used an olfactory starvation-conditioning paradigm that probes defined chemosensory and interneuron circuits and provides quantitative readouts of learning and short-term memory in early adulthood (34, 55–57). Synchronized Day 5 adults (cultured in the presence of FUDR) were evenly divided onto three unseeded 60 mm NGM plates: naïve, conditioning-1, and conditioning-2. For conditioned groups, 10 µL pure isoamyl alcohol (IAA; Sigma-Aldrich W205508) was applied to the plate lid. Plates were sealed with Parafilm, inverted, and incubated at room temperature for 90 min. Naïve animals were handled in parallel without IAA.

After conditioning, animals were washed in M9 and transferred to chemotaxis plates (unseeded 100 mm NGM plates) pre-spotted with 1 M sodium azide (Sigma-Aldrich S2002) at each scoring locus. See Fig. 4a for a schematic. Two attractant spots received 4 µL of 1:50 IAA in ultrapure water, and two control spots received 4 µL ultrapure water. Animals were allowed to undergo chemotaxis for 2 h at room temperature. Naïve and conditioning-1 groups were tested immediately after conditioning to assess learning. Conditioning-2 animals were maintained on unseeded plates for 1 h after conditioning, then tested to assess short-term memory.

Chemotaxis Index (CI) = (N_IAA − N_control) / N_total. Learning Index (LI) = CI_naive − CI_trained, where “trained” refers to conditioning-1 for learning at 0 h post-conditioning and conditioning-2 for memory at 1 h post-conditioning. Each RNAi condition included three to four independent replicates with at least 100 animals per replicate per condition.

### Generation of Transgenes

*tbc-17* overexpression constructs were generated using Gateway cloning (Thermo Fisher Scientific). All constructs were verified by Sanger sequencing prior to use. Transgenic animals were produced by gonadal microinjection, with plasmid DNA at 30 ng/µL (PELZ138: *tbc-17p::tbc-17* cDNA) and the co-injection marker *ttx-3p::GFP* at 50 ng/µL into the *[zdIs5(mec-4p::GFP)*, *uIs69(myo-2p::mCherry + unc-119p::sid-1)]* background, as detailed in Table S4. Two independent transgenic lines were assayed.

### Gentle Touch Assay

Mechanosensory responses were assessed on thinly seeded NGM plates at room temperature following established protocols (105). Animals acclimated for 5 minutes before testing. Touch stimuli were delivered with an eyelash mounted on a toothpick. Five gentle strokes were applied along the lateral side of the posterior third of the body. A response was scored positive if the animal initiated forward locomotion. The mechanosensory-defective strain TU253 *[mec-4(u253)]* served as the touch-insensitive control. Assays were performed on wild-type Day 3 and Day 9 animals, and on *tbc-17* overexpression animals at Day 3, which were cultured in the presence of FUDR.

### Neuronal Mitochondrial Morphology Analysis and Heat Stress Assay

Mitochondrial morphology in PLM neurons was visualized using the *jsIs609 [mec-4p::mitoGFP]* reporter in the *uIs69 (myo-2p::mCherry + unc-119p::sid-1)* background. Animals received lifelong *tbc-17* RNAi or EV treatment in the absence of FUDR. Animals were mounted on 4% agarose pads and anesthetized with 5 mM levamisole (Sigma-Aldrich, Cat# L9756) in M9 buffer. Imaging was performed on a Zeiss Axio Imager M2 using a 63× oil-immersion objective. Z-stacks were acquired at 1 µm per slice, and exposure settings were kept constant across conditions. Z-stacks were processed as XY maximum-intensity projections. Mitochondrial features were quantified in a defined 220 µm segment at the distal end of the PLM neurite in each animal (68). For each animal, we calculated four metrics: mean mitochondrial particle area, mean aspect ratio (major axis divided by minor axis for each mitochondrial particle), load (cumulative mitochondrial area per 100 µm of neurite length), and density (number of mitochondrial particles within the distal 220 µm of the neurite). Image analysis was performed in FIJI (NIH) using thresholding and the Analyze Particles function.

For heat stress induction, animals were incubated at 37 °C for 1 hour on RNAi plates seeded with HT115(DE3) bacteria carrying the empty L4440 vector (control) or *tbc-17* RNAi, allowed to recover for 4 hours at 20 °C, and then imaged. Each condition included at least 20 animals across three biological replicates.

### Amyloid-beta-driven Neurodegeneration Assay

To assess Aβ-induced neurotoxicity in PVD neurons, the transgenic strain *gnaIs2 [myo-2p::YFP + unc-119p::Abeta1–42]* or its control strain *gnaIs1 [myo-2p::yfp]* (31) was crossed with the PVD reporter *wdIs51 [F49H12.4p::GFP]*, with or without *uIs69 [unc-119p::sid-1]*. A transgenic strain *dvIs14 [(pCL12) unc-54::beta 1-42 + (pCL26) mtl-2::GFP]* which expresses human Aβ specifically in muscles or its control strain *dvIs15 [(pPD30.38) unc-54(vector) + (pCL26) mtl-2::GFP]* was also crossed with another PVD reporter *lxyEx83 [ser-2*(*3*)*p::GFP-utRCH, ser-2*(*3*)*p::mCherry-pH].* PVD dendritic beading was scored using established criteria as above. Experiments were performed in the presence of FUDR.

### RT-qPCR

To assess RNAi knockdown efficiency and gene expression levels by RT-qPCR, we used a modified 10-worm lysis protocol (106). Ten worms were rinsed in ultrapure water and transferred to 1 µL of lysis buffer (0.25 mM EDTA, 5 mM Tris pH 8.0, 0.5% Triton X-100, 0.5% Tween-20, 1 mg/mL proteinase K) in PCR tubes. Samples were incubated at 65 °C for 15 min, then at 85 °C for 1 min, and stored at −80 °C. After thawing, 1.5 µL of fresh lysis buffer was added and the thermal cycle was repeated. Lysates were treated with dsDNase (Thermo Fisher, Cat# EN0771) prior to reverse transcription. cDNA was synthesized with iScript Reverse Transcription Supermix (Bio-Rad, Cat# 1708841). qPCR was performed using SYBR Green (Bio-Rad, Cat# 1725120) or gene-specific TaqMan assays (Thermo Fisher) on a CFX96 Real-Time PCR System (Bio-Rad). Relative expression was calculated by the ΔΔCT method, normalized to *cdc-42*, *ama-1*, or *act-2*. Representative results for RNAi efficiency are shown in Fig. S1.

### Transcriptomic Datasets and Analysis

Gene expression datasets (GSE118553, GSE44770, GSE48350, GSE53890 and GSE176088) were obtained from the NCBI Gene Expression Omnibus. Data were analyzed using GEO2R to identify differentially expressed genes (DEGs), with criteria of a *p*□<□0.05.

### Statistical Analysis

All analyses were performed using GraphPad Prism (v10.4.0). Normality was assessed using the Shapiro–Wilk test. Survival data were evaluated using Kaplan–Meier analysis with log-rank (Mantel–Cox) tests. For multiple group comparisons, one-way ANOVA with Tukey’s post hoc test was used for normally distributed datasets, and Kruskal–Wallis tests with Dunn’s correction were used for non-parametric datasets. For two group comparisons, unpaired two-tailed *t*-tests for parametric data or Mann–Whitney tests for non-parametric data were used. Fisher’s exact test was also used where applicable. Severity for PVD dendritic beading and PLM ectopic branching were analyzed using the Cochran–Mantel–Haenszel trend test across independent experiments, stratified by replicate (107). Two-way ANOVA with post hoc testing was used for mitochondrial morphology analyses under basal and heat stress conditions. Significance was defined as *p* < 0.05. Results are reported as mean ± standard error of the mean (SEM), unless otherwise specified. Sample sizes and statistical tests are provided in the corresponding figure legends.

