## Supplementary Information for "Functional Analysis of Late-Onset Alzheimer’s Disease Risk Genes in *Caenorhabditis elegans* Identifies Regulators of Neuronal Aging"

**Figure S1. *C. elegans* LOAD gene homolog expression during aging and RNAi efficiency.**

**a** mRNA expression changes of LOAD gene homologs in Day 10 animals compared with L4, based on the GSE176088 dataset.

**b-c** Relative mRNA levels of *ech-2* following *ech-2* RNAi treatment (Day 3) and *T09B9.4* following *T09B9.4* RNAi treatment (Day 14), compared with EV. Expression values were normalized to *ama-1*, *cdc-42*, or *act-2* as housekeeping genes.

Statistical comparisons were performed using Mann–Whitney tests. All experiments were conducted in the presence of FUDR. EV, control animals fed with the empty L4440 RNAi vector. **$p < 0.01$; ns, not significant.

**Figure S2. PVD dendritic beading following lifelong RNAi knockdown of LOAD-associated gene homologs.**

**a** Quantification of dendritic beading severity in Day 3 PVD neurons following lifelong RNAi knockdown of selected LOAD gene homologs.

**b** Quantification of dendritic beading severity in Day 11 PVD neurons following lifelong RNAi knockdown of selected LOAD gene homologs.

PVD beading severity was scored in four categories (Normal, Mild, Moderate, Severe) based on bead number across the dendritic tree and assigned ordinal values 0–3 (see Methods). For each RNAi condition and replicate, a Δ-Severity value was calculated as the average severity score in the LOAD gene RNAi group minus that of its paired EV control to visualize the direction and magnitude of morphological change; Δ-Severity was used only for descriptive plotting.

**Figure S3. Knockdown of LOAD gene homologs does not affect the aging-associated sharp bends/kinks in PLM neurite, or overall PLM neuritic structure in young adults.**

**a–f** Quantification of sharp bends or kinks in Day 3 PLM neurons following lifelong RNAi knockdown of LOAD gene homologs. The number of sharp bends or kinks was quantified along the main PLM sensory dendrite. Data represent one of the three independent biological replicates ($n \geq 30$ animals per condition). Statistical comparisons were performed using Fisher's exact test.

**g–l** Quantification of sharp bends or kinks in Day 9 PLM neurons following lifelong RNAi knockdown of LOAD gene homologs. The number of sharp bends or kinks was quantified along the main PLM sensory dendrite. Data represent one of the three independent biological replicates ($n \geq 30$ animals per condition). Statistical comparisons were performed using Fisher's exact test.

**m–r** Quantification of ectopic branching severity in Day 3 PLM neurons following lifelong RNAi knockdown of LOAD gene homologs. Ectopic branching was scored on a four-point ordinal scale based on the number of ectopic branches (0 = normal, 1 = mild, 2 = moderate, 3+ = severe). Data represent one of the three independent biological replicates ($n \geq 30$ animals per condition). Statistical comparisons were performed using Fisher's exact test.

All experiments were performed in the presence of FUDR. EV indicates control animals fed with the empty L4440 RNAi vector. ns indicates not significant.

**Figure S4. Neuronal expression patterns of *R166.2* vs *C36B7.6*.**

**a** Δ% expressing by cell type. Difference in penetrance per cell type, defined as % cells expressing *R166.2* − % cells expressing *C36B7.6*. Positive values indicate more cells expressing *R166.2*.

**b** Δ Transcripts Per Million (TPM) by cell type. Positive values indicate more expression of *R166.2* in corresponding cell type.

Source data: WormBase Expression tables (CeNGEN (1)). Cell type names follow WormBase annotations.

**Figure S5. Associative learning and memory-like behavior in *C. elegans*.**

**a–b** Quantification of chemotaxis behavior following conditioning in young (Day 1) and mid-adult (Day 5) animals. Data are shown for trained versus naïve groups at 0 h post-conditioning (PC; learning acquisition, a) and 1 h post-conditioning (PC; short-term memory retention, b).

**c–h** Quantification of naive chemotaxis at Day 5 following lifelong RNAi knockdown of selected LOAD gene homologs (n = 100 animals per condition).

Each condition represents pooled results from at least three independent biological replicates (n = 100 animals per condition). Statistical comparisons were performed using unpaired two-tailed t-tests. All experiments were performed in the presence of FUDR. EV, control animals fed with the empty L4440 RNAi vector. ns, not significant.

**Figure S6. Characterization of *tbc-17*.**

**a** Clustal-format sequence alignment of TBC-17 and human USP6NL generated with MAFFT (v7.511). Invariant, conserved, and semi-conserved residues are indicated by an asterisk (*), colon (:), and period (.), respectively.

**b** Ectopic branching severity in young (Day 3) and old (Day 9) PLM neurons following lifelong *tbc-17* RNAi using a second RNAi clone (PELZ136). Data represent pooled results from independent biological replicates (n ≥ 90 animals per condition). Experiments with FUDR.

**c** Lifespan of animals subjected to adulthood-specific *tbc-17* RNAi compared with EV. Experiments performed with CZ10175 animals without neuronal RNAi enhancement. Experiments performed with FUDR.

**d** Lifespan of *tbc-17* overexpression (ELZ279, *tbc-17* OE) compared with control animals (ELZ238). Experiments without FUDR.

**e** Ectopic branching quantification in Day 3 PLM neurons in control (CT, ELZ238) (n = 169) and *tbc-17* OE animals (ELZ278, line 2) (n = 130) fed with *E. coli* OP50. Experiments with FUDR.

**f** Relative *tbc-17* mRNA levels at L4-stage in *tbc-17* OE animals (ELZ279), compared with control animals (ELZ238). Expression was normalized to *cdc-42* or *act-2*. L4 animals were used for this analysis.

**g** Gentle touch response in control animals (ELZ238) at Day 3 (young, n = 10) and Day 9 (old, n = 30). Experiments with FUDR.

**h** Gentle touch response in control animals (ELZ238, n = 20) and *tbc-17* OE animals (ELZ279, n = 20) at Day 3. Experiments with FUDR.

**i–l** Mitochondrial morphology in PLM neurons EV and *tbc-17* RNAi animals at Day 2 and Day 5 under basal conditions. Metrics include area (i), shape (j), load (k), and density (l). Experiments without FUDR.

**m–n** Relative *drp-1* (m) and *fzo-1* (n) mRNA levels at Day 5 following *tbc-17* RNAi compared with EV controls, normalized to *cdc-42* or *act-2.* Experiments with FUDR.

Statistical tests: (b) stratified CMH; (c–d) log-rank; (e) Fisher's exact; (f–h, m–n) Mann–Whitney; (i–l) two-way ANOVA. EV, empty L4440 RNAi vector. Significance: **$p < 0.01$, ***$p < 0.001$, ****$p < 0.0001$; ns, not significant.

**Figure S7.** ***ech-2*****'s effect on Aβ-induced PVD dendritic beading.**

**a** Quantification of dendritic beading in Day 1 PVD neurons of control animals (Ctrl strain - ELZ273, *uIs69; wdIs51; gnaIs1*, n = 37) and Aβ overexpression animals (Aβ OE - ELZ266, *uIs69; wdIs51; gnaIs2*, n = 36).

**b** Quantification of dendritic beading severity in old (Day 9) PVD neurons following RNAi knockdown of *ech-2* in the Aβ-overexpressing background. Control strain (ELZ273, *uIs69; wdIs51; gnaIs1*) with EV, n = 23); Aβ OE strain (ELZ266, *uIs69; wdIs51; gnaIs2*) with EV (n = 34) or with *ech-2* RNAi (n = 25).

**c** Quantification of dendritic beading in PVD neurons at Day 3 for control (CT) animals (ELZ179, *wdIs51*; *gnaIs1*, n = 30) and animals overexpressing Aβ (ELZ186, *wdIs51*; *gnaIs2*, n = 29), and at Day 7 for control animals (n = 35) and Aβ overexpression animals (n = 35). Without neuronal RNAi-enhanced background.

**d** Quantification of dendritic beading in Day 1 PVD neurons of control (CT) animals (ELZ214, *dvIs15; lxyEx83*, n = 12) and animals expressing human Aβ in muscle tissues (ELZ199, *dvIs14; lxyEx83*, n = 16). Without neuronal RNAi-enhanced background.

**e** Relative mRNA levels of *nlp-29* after *ech-2* RNAi treatment compared with EV. *cdc-42* as a housekeeping gene.

Statistical comparisons were performed using Fisher’s exact test (a–d) and Mann–Whitney tests (e). All experiments were conducted in the presence of FUDR. $^{**}p < 0.01$; ns, not significant.

Table S1: Human brain transcriptomic data relevant to Table 1.

Table S2: Lifespan data

Table S3: PVD dendritic beading data

Table S4: Strain information

Table S5: Plasmid information

Table S6: Primer information

#### Figure S1

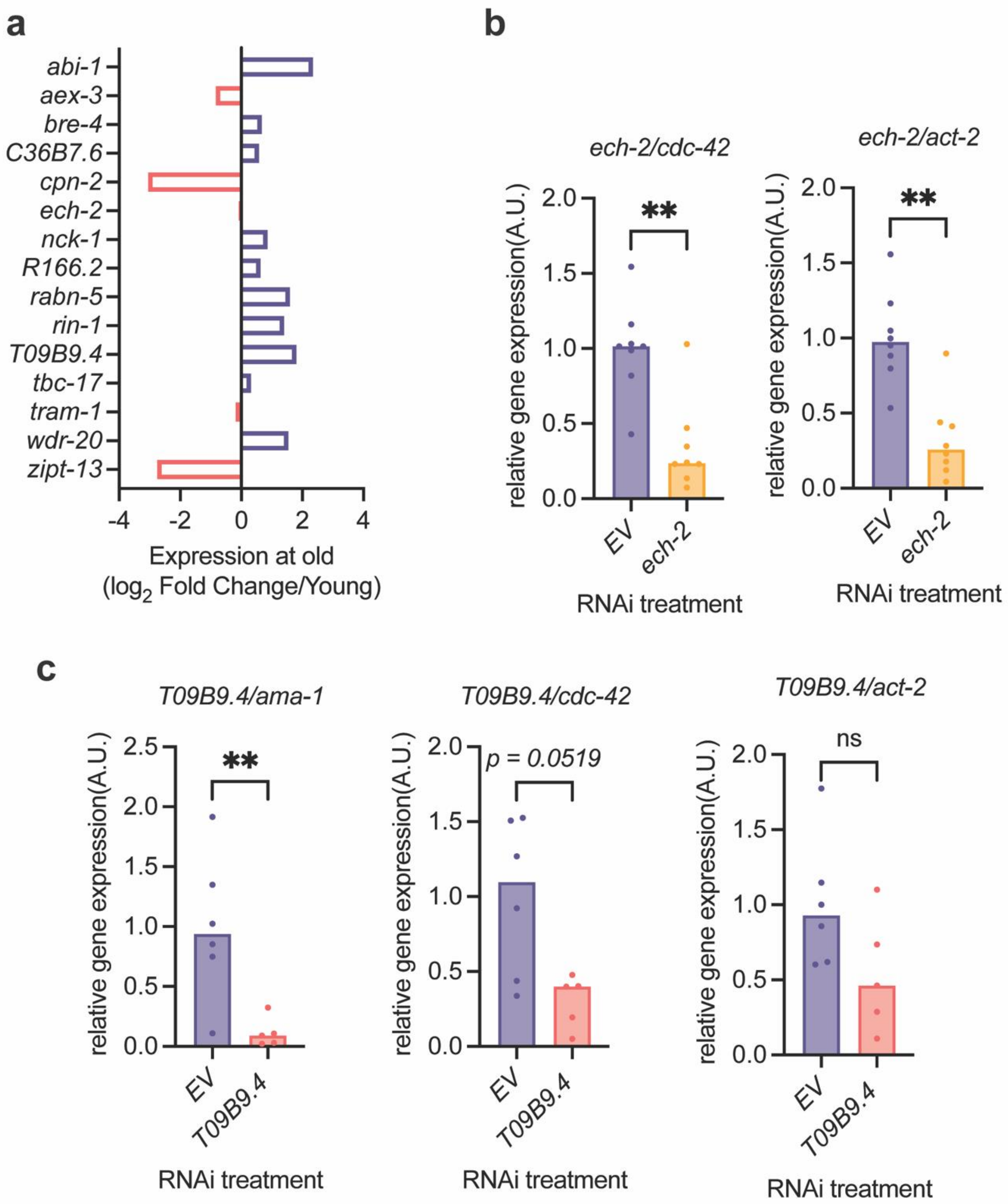

a
abi-1
aex-3
bre-4
C36B7.6
cpn-2
ech-2
nck-1
R166.2
rabn-5
rin-1
T09B9.4
tbc-17
tram-1
wdr-20
zipt-13
-4
-2
0
2
4
Expression at old
(log2 Fold Change/Young)
b
ech-2/cdc-42
ech-2/act-2
relative gene expression(A.U.)
**
EV
ech-2
RNAi treatment
c
T09B9.4/ama-1
T09B9.4/cdc-42
T09B9.4/act-2
p = 0.0519
ns
T09B9.4
RNAi treatment

### Figure S2

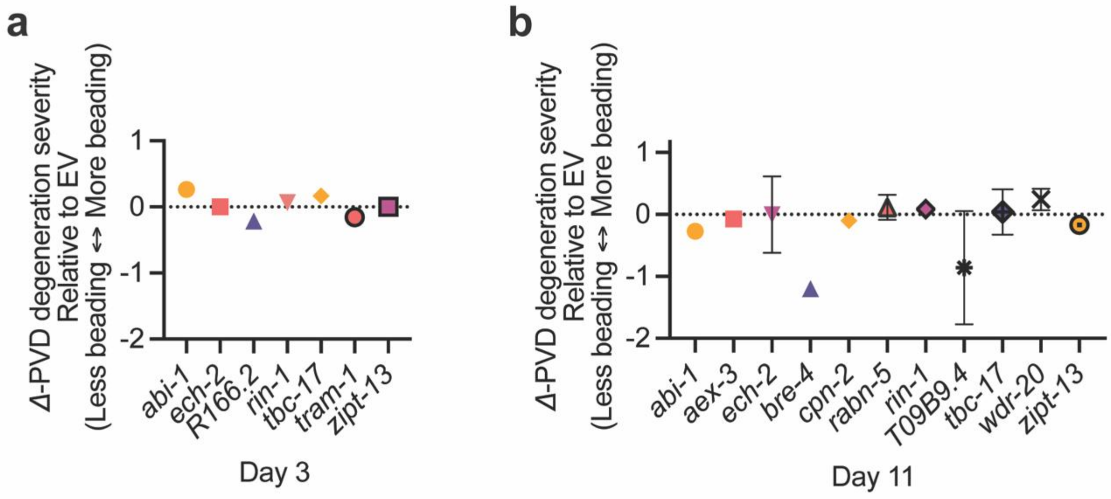
a
Δ-PVD degeneration severity
Relative to EV
(Less beading ↔ More beading)
1
0
-1
-2
abi-1
ech-2
R166.2
rin-1
tbc-17
tram-1
zipt-13
Day 3
b
Δ-PVD degeneration severity
Relative to EV
(Less beading ↔ More beading)
1
0
-1
-2
abi-1
aex-3
ech-2
bre-4
cpn-2
rabn-5
rin-1
T09B9.4
tbc-17
wdr-20
zipt-13
Day 11

### Figure S3

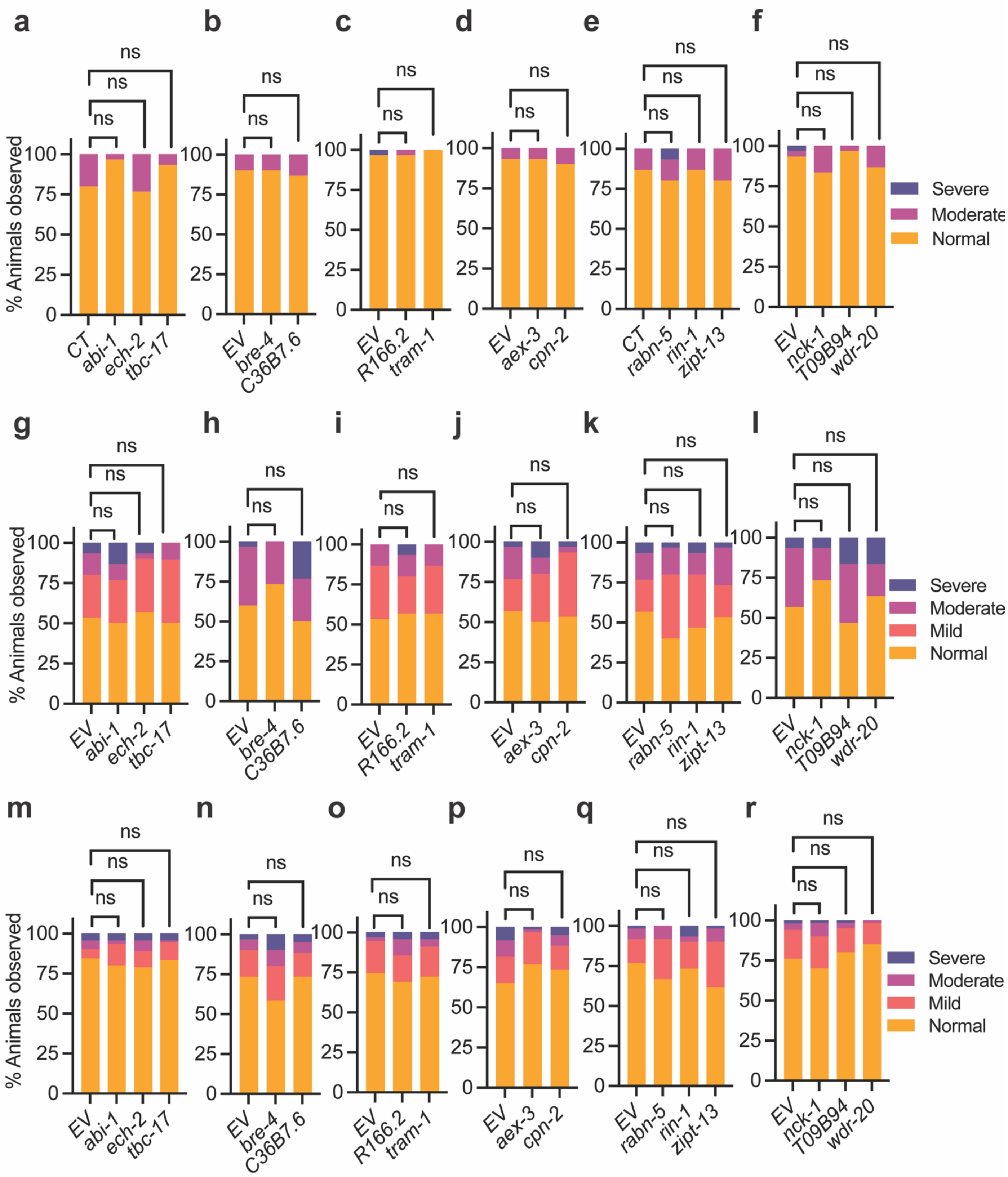

a
b
c
d
e
f
g
h
i
j
k
l
m
n
o
p
q
r
ns
% Animals observed
0
25
50
75
100
CT
abi-1
ech-2
tbc-17
EV
bre-4
C36B7.6
R166.2
tram-1
aex-3
cpn-2
rabn-5
rin-1
zipt-13
nck-1
T09B94
wdr-20
Severe
Moderate
Mild
Normal

#### Figure S4

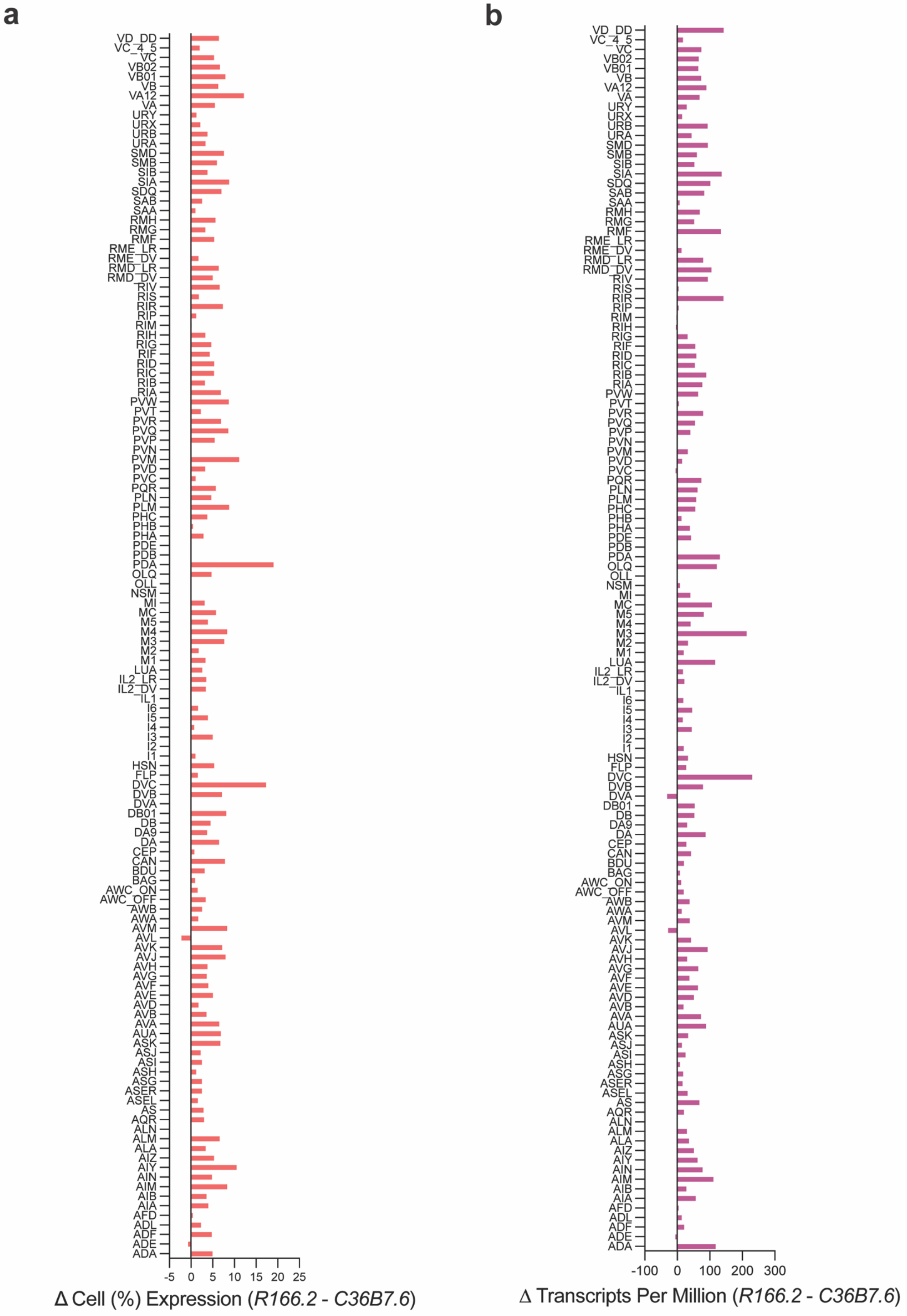
a
VD_DD
VC_4_5
VC
VB02
VB01
VB
VA12
VA
URY
URX
URB
URA
SMD
SMB
SIB
SIA
SDQ
SAB
SAA
RMH
RMG
RMF
RME_LR
RME_DV
RMD_LR
RMD_DV
RIV
RIS
RIR
RIP
RIM
RIH
RIG
RIF
RID
RIC
RIB
RIA
PVW
PVT
PVR
PVQ
PVP
PVN
PVM
PVD
PVC
PQR
PLN
PLM
PHC
PHB
PHA
PDE
PDB
PDA
OLQ
OLL
NSM
MI
MC
M5
M4
M3
M2
M1
LUA
IL2_LR
IL2_DV
IL1
I6
I5
I4
I3
I2
I1
HSN
FLP
DVC
DVB
DVA
DB01
DB
DA9
DA
CEP
CAN
BDU
BAG
AWC_ON
AWC_OFF
AWB
AWA
AVM
AVL
AVK
AVJ
AVH
AVG
AVF
AVE
AVD
AVB
AVA
AUA
ASK
ASJ
ASI
ASH
ASG
ASER
ASEL
AS
AQR
ALN
ALM
ALA
AIZ
AIY
AIN
AIM
AIB
AIA
AFD
ADL
ADF
ADE
ADA
-5 0 5 10 15 20 25
Δ Cell (%) Expression (*R166.2 - C36B7.6*)
b
VD_DD
VC_4_5
VC
VB02
VB01
VB
VA12
VA
URY
URX
URB
URA
SMD
SMB
SIB
SIA
SDQ
SAB
SAA
RMH
RMG
RMF
RME_LR
RME_DV
RMD_LR
RMD_DV
RIV
RIS
RIR
RIP
RIM
RIH
RIG
RIF
RID
RIC
RIB
RIA
PVW
PVT
PVR
PVQ
PVP
PVN
PVM
PVD
PVC
PQR
PLN
PLM
PHC
PHB
PHA
PDE
PDB
PDA
OLQ
OLL
NSM
MI
MC
M5
M4
M3
M2
M1
LUA
IL2_LR
IL2_DV
IL1
I6
I5
I4
I3
I2
I1
HSN
FLP
DVC
DVB
DVA
DB01
DB
DA9
DA
CEP
CAN
BDU
BAG
AWC_ON
AWC_OFF
AWB
AWA
AVM
AVL
AVK
AVJ
AVH
AVG
AVF
AVE
AVD
AVB
AVA
AUA
ASK
ASJ
ASI
ASH
ASG
ASER
ASEL
AS
AQR
ALN
ALM
ALA
AIZ
AIY
AIN
AIM
AIB
AIA
AFD
ADL
ADF
ADE
ADA
-100 0 100 200 300
Δ Transcripts Per Million (*R166.2 - C36B7.6*)

#### Figure S5

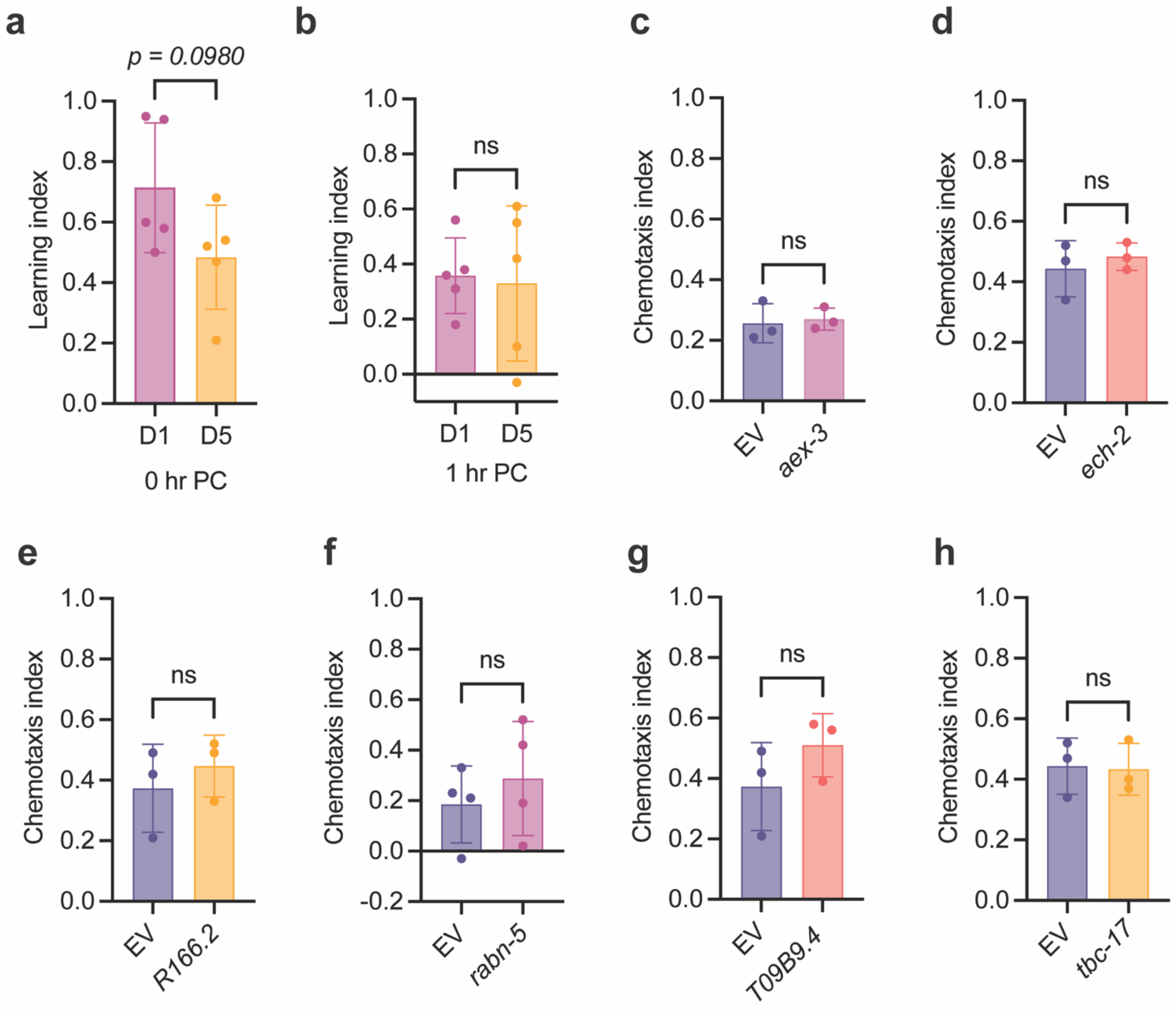

a
p = 0.0980
Learning index
1.0
0.8
0.6
0.4
0.2
0.0
D1
D5
0 hr PC
b
Learning index
ns
D1
D5
1 hr PC
c
Chemotaxis index
ns
EV
aex-3
d
Chemotaxis index
ns
EV
ech-2
e
Chemotaxis index
ns
EV
R166.2
f
Chemotaxis index
ns
-0.2
EV
rabn-5
g
Chemotaxis index
ns
EV
T09B9.4
h
Chemotaxis index
ns
EV
tbc-17

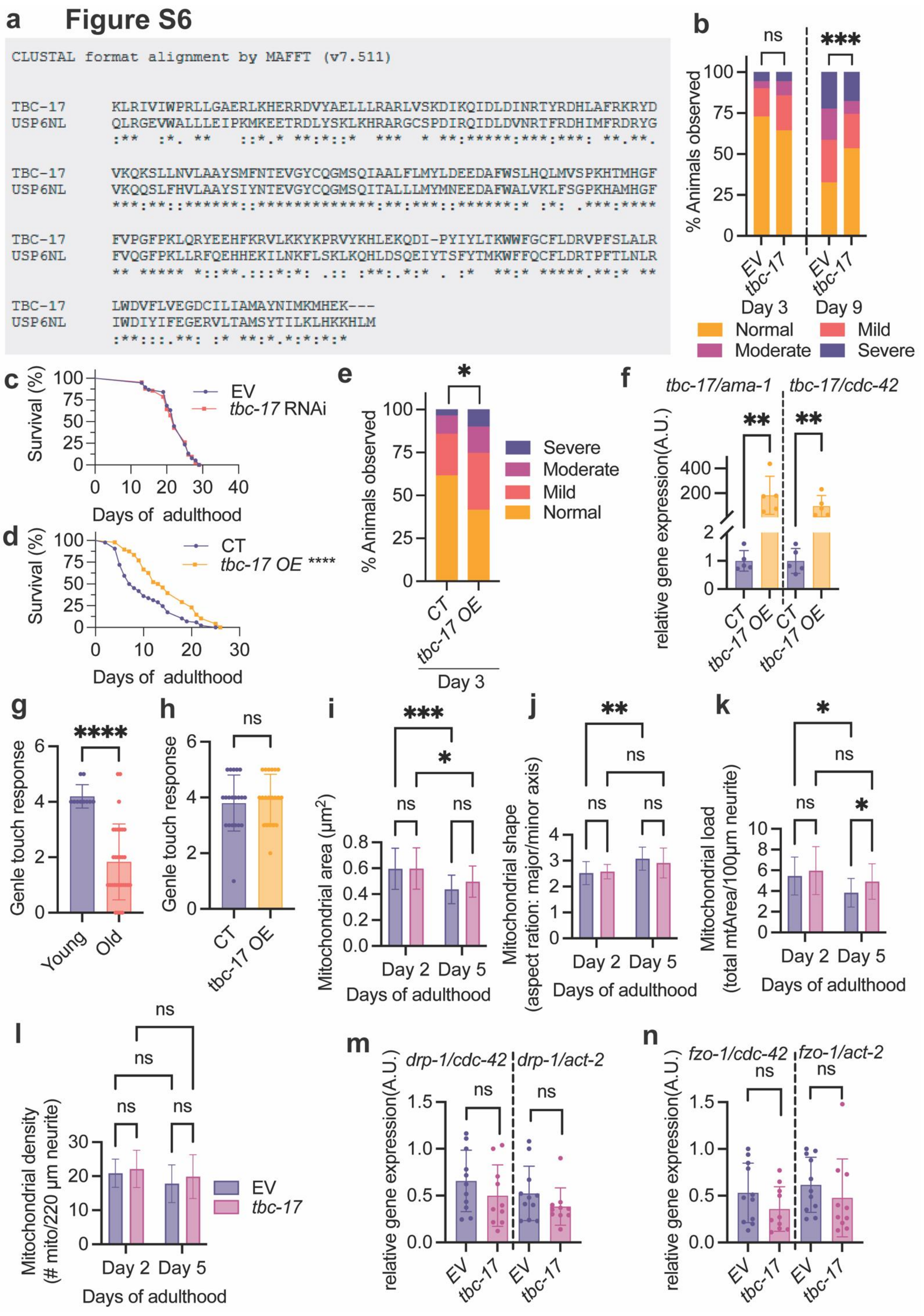
a
Figure S6
CLUSTAL format alignment by MAFFT (v7.511)
TBC-17 KLRIVIWPRLLGAERLKHERRDVYAELLLRARLVSKDIKQIDLDINRTYRDHLAFRKRYD
USP6NL QLRGEVWALLLEIPKMKEETRDLYSKLKHRARGCSPDIRQIDLDVNRTFRDHIMFRDRYG
:** :*. ** ::*.* **:*::* *** * **:*****:***:***: **.**.
TBC-17 VKQKSLLNVLAAYSMFNTEVGYCQGMSQIAALFLMYLDEEDAFWSLHQLMVSPKHTMHGF
USP6NL VKQQSLFHVLAAYSIYNTEVGYCQGMSQITALLLMYMNEEDAFWALVKLFSGPKHAMHGF
***:**::******::*************:**:***::******:* :*: .***:****
TBC-17 FVPGFPKLQRYEEHFKRVLKKYKPRVYKHLEKQDI-PYIYLTKWWFGCFLDRVPFSLALR
USP6NL FVQGFPKLLRFQEHHEKILNKFLSKLKQHLDSQEIYTSFYTMKWFFQCFLDRTPFTLNLR
** ***** *::**.:::*:*: .:: :**:.*:* . :* **:* *****.**:* **
TBC-17 LWDVFLVEGDCILIAMAYNIMKMHEK---
USP6NL IWDIYIFEGERVLTAMSYTILKLHKKHLM
:**:::.**: :* **:*.*:*:*:*
b
ns
***
% Animals observed
EV
tbc-17
Day 3
Day 9
Normal
Mild
Moderate
Severe
c
Survival (%)
EV
tbc-17 RNAi
Days of adulthood
d
Survival (%)
CT
tbc-17 OE ****
Days of adulthood
e
*
% Animals observed
Severe
Moderate
Mild
Normal
CT
tbc-17 OE
Day 3
f
tbc-17/ama-1
tbc-17/cdc-42
**
relative gene expression(A.U.)
CT
tbc-17 OE
g
****
Genle touch response
Young
Old
h
ns
Genle touch response
CT
tbc-17 OE
i
***
*
ns
Mitochondrial area (μm²)
Day 2
Day 5
Days of adulthood
j
**
ns
Mitochondrial shape (aspect ration: major/minor axis)
Day 2
Day 5
Days of adulthood
k
*
ns
Mitochondrial load (total mtArea/100μm neurite)
Day 2
Day 5
Days of adulthood
l
ns
Mitochondrial density (# mito/220 μm neurite)
EV
tbc-17
Day 2
Day 5
Days of adulthood
m
drp-1/cdc-42
drp-1/act-2
ns
relative gene expression(A.U.)
EV
tbc-17
n
fzo-1/cdc-42
fzo-1/act-2
ns
relative gene expression(A.U.)
EV
tbc-17

### Figure S7

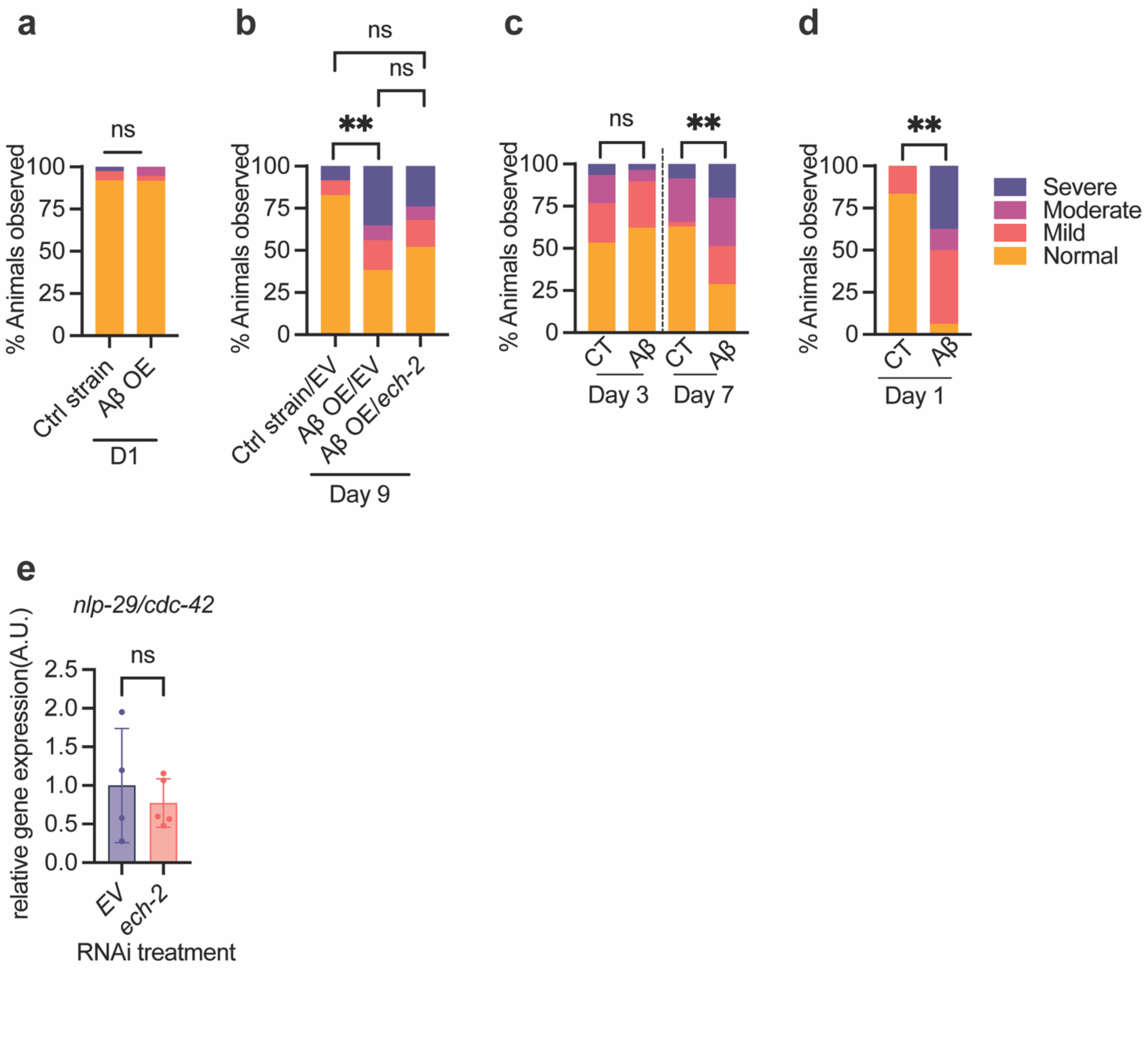

a
ns
% Animals observed
100
75
50
25
0
Ctrl strain
Aβ OE
D1
b
ns
ns
**
% Animals observed
Ctrl strain/EV
Aβ OE/EV
Aβ OE/ech-2
Day 9
c
ns
**
% Animals observed
CT
Aβ
CT
Aβ
Day 3
Day 7
d
**
% Animals observed
CT
Aβ
Day 1
Severe
Moderate
Mild
Normal
e
nlp-29/cdc-42
ns
relative gene expression(A.U.)
2.5
2.0
1.5
1.0
0.5
0.0
EV
ech-2
RNAi treatment
